# Listeria motility increases the efficiency of goblet cell invasion during intestinal infection

**DOI:** 10.1101/2022.07.21.500464

**Authors:** Inge M. N. Wortel, Seonyoung Kim, Annie Y. Liu, Enid C. Ibarra, Mark J. Miller

**Author notes:** These authors contributed equally to the work (Inge M. N. Wortel, Seonyoung Kim).

## Abstract

*Listeria monocytogenes* (Lm) is a food-borne pathogen that causes severe bacterial gastroenteritis, with high rates of hospitalization and mortality. Lm is ubiquitous in soil, water and livestock, and can survive and proliferate at low temperatures. Following oral ingestion of contaminated food, Lm crosses the epithelial through intestinal goblet cells in a mechanism depending on Lm InlA and host E-cadherin. Importantly, human infections typically occur with Lm growing at or below room temperature, which are flagellated and motile. Even though many important human bacterial pathogens are flagellated, little is known regarding the effect of bacterial motility on invasion and immune evasion.

Here, we used complementary imaging and computer modeling approaches to test the hypothesis that bacterial motility helps Lm locate and engage target cells permissive for invasion. Imaging explanted mouse and human intestine, we confirmed that Lm grown at room temperature uses motility to scan the epithelial surface and preferentially attach to target cells. Furthermore, we integrated quantitative parameters from our imaging experiments to construct a versatile “layered” cellular Potts model (L-CPM) that simulates host-pathogen dynamics. Simulated data are consistent with the hypothesis that bacterial motility enhances invasion by allowing bacteria to search the epithelial surface for their preferred invasion targets. Indeed, our model consistently predicts that motile bacteria have invaded ∼2-fold more at the 1-hour mark. This invasion advantage persists even in the presence of host phagocytes, with the balance between invasion and phagocytosis governed almost entirely by bacterial motility.

In conclusion, our simulations provide insight into host pathogen interactions and challenge fundamental assumptions regarding how phagocytes might limit bacterial invasion early during infection.

## Introduction

Gastroenteritis and diarrheal diseases are a major source of mortality and morbidity worldwide, especially among children in developing countries ^1,2^. The mucosal barrier of the intestine is exposed continuously to microbes and has evolved a variety of mechanisms to prevent pathogen invasion, including epithelial cell shedding, secretion of antimicrobial peptides and mucus and mucosal antibody production ^3–5^. To counter these host defense mechanisms, enteropathogenic bacteria produce virulence factors to promote barrier breach, host cell invasion and inhibit the immune response ^3^. Understanding the early stages of bacterial pathogenesis is important for identifying vaccine targets and developing new therapies for bacterial infections.

*Listeria monocytogenes* (Lm) is a food borne pathogen that causes severe life-threatening disease with high hospitalization and mortality rates ^6^. Lm is ubiquitous in soil, water and livestock and can survive and proliferate at low temperatures, for example in refrigerated foods such as deli meats, unpasteurized soft cheeses, and smoked fish ^7^. Lm expresses a variety of virulence factors ^8,9^ that facilitate intracellular invasion and the evasion of host immune responses. One example is the forming protein Listeriolysin O that disrupts the phagolysosome membrane, allowing Lm to escape into the cytosol and replicate intracellularly. Another is ActA, which polymerizes host cell actin to propel Lm directly into neighboring cells to infect them ^10^, evading humoral and innate immunity.

Lm infection has been widely studied using laboratory mouse models and i.v. infection routes ^11^. However, efficient oral infection requires the binding of Lm InlA to E-cadherin, which is human-specific ^11,12^. To overcome this limitation, “murinized” Lm was engineered to express a mutated form of InlA that binds mouse E-cadherin and allows oral infection ^13–15^. Alternatively, transgenic mice expressing human E-cadherin can be infected orally with WT (wild-type) Lm ^16^ to model physiological Lm invasion ^17^. Such studies have demonstrated that on the luminal side of the gut epithelium, E-cadherin is primarily accessible around goblet cells, a subset of secretory epithelial cells. Thus, goblet cells are the preferred target cell for attachment and transcytosis across the epithelial barrier ^12,16,18,19^. However, the key system parameters and cell dynamics that lead to these Lm-goblet cell interactions are less well understood.

In contrast to published studies using mouse models, human Lm infections typically occur after the ingestion of contaminated food with bacteria proliferating at or below room temperature (RT, 20-25°C), which are typically flagellated and motile ^7,20–22^. Indeed, many clinically important bacterial pathogens are flagellated and motile, including Salmonella, Campylobacter, Helicobacter, Yersinia, Pseudomonas ^23–25^. This raises the question: to what degree does bacterial motility determine immune outcomes? Previous studies have been contradictory: one study found that Lm flagellar null mutants show similar immunological outcomes in mice and that deletion of the *flaA* gene repressor *mogR* dramatically decreased virulence in vivo ^20^, while others showed that flagellated Lm have a competitive advantage early during oral infection ^22^.

Here, we investigated how bacterial motility impacts target cell engagement and invasion efficiency. Two-photon (2P) imaging of explanted mouse and human intestine showed that motile Lm explored the surface of the epithelium and rapidly accumulated around target cells. To predict the consequences of Lm motility for Lm-goblet cell interactions and Lm invasion over time, under various environmental and bacterial motility scenarios, we constructed a computational model called the cellular Potts model (CPM) ^26–28^. Using interacting layers of epithelial cells, motile bacteria, and (potentially) host phagocytes, our “layered CPM” (L-CPM) integrates quantitative parameters obtained from in vivo, in vitro and explant experiments to simulate host-pathogen dynamics at the epithelium. This model reveals that bacterial motility enhances invasion efficiency by permitting bacteria to rapidly search for and engage permissive target cells. This motility advantage remains even in the presence of (some) host phagocytes – but disappears in conditions where phagocytosis is so dominant that bacteria are phagocytosed before they can invade. Surprisingly, we predict that this balance between invasion and phagocytosis is governed almost entirely by bacterial motility, with only a minor role for phagocyte motility. In conclusion, our simulations provide insight into host pathogen interactions and challenge fundamental assumptions regarding how phagocytes might limit bacterial invasion early during infection.

## Results

### Murinized Lm cultured at RT is flagellated and highly motile

First, we evaluated the effect of culture temperature on flagella formation and motility of murinized Listeria. Lm was cultured in Brain Heart Infusion media (BHI) with shaking at room temperature (Lm-RT) or at 37°C (Lm-37), and flagella formation was assessed by transmission electron microscopy. Lm-37 (Fig. 1A,B) were non-flagellated while Lm-RT showed multiple long peritrichous flagella, similar to published findings with the wild type strain EGD Lm^Wt 22^, confirming that RT culture produces flagellated murinized Lm bacteria.

**Figure 1:**
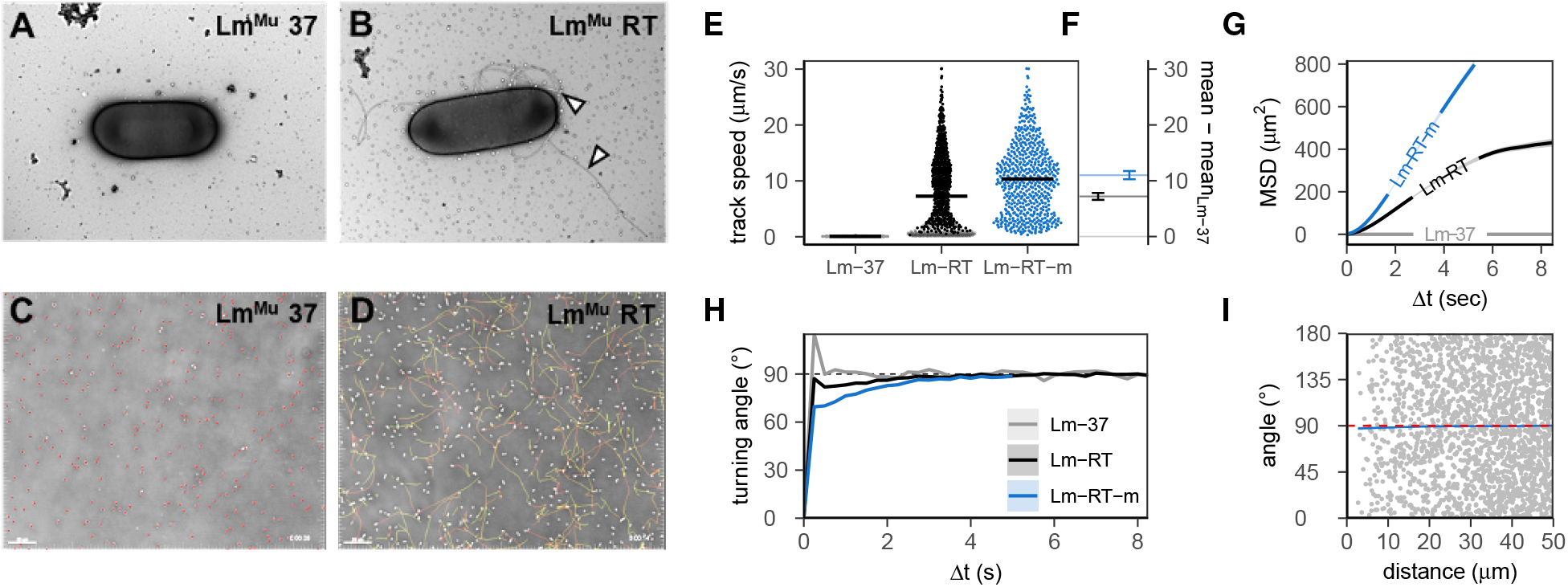
Murinized Listeria (Lm^Mu^) grown at room temperature are flagellated and highly motile. **A** electron microscopy shows that Lm cultured at 37°C (Lm-37) are non-flagellated, while **B**, Lm grown at 23°C (Lm-RT) develop extensive peritrichous flagella (white arrows). Scale bar=1μm. **C**, video microscopy shows that Lm-37 are non-motile in vitro (white cells, time encoded tracks) in contrast to **D**, Lm-RT which are highly motile (white cells, time encoded tracks). Scale bar=30 μm. This difference in motility is supported by **E**, the mean speed per bacterium with Lm-RT-m a filtered, “motile” subset; see Supp. Fig. S1), and **F**, the bootstrapped 99% confidence interval (CI) of the increase in population mean speed for Lm-RT(-m) versus Lm-37 (bootstrap N = 10000). The increased motility of Lm-RT(-m) also yields higher **G**, mean squared displacements (MSD ± standard error, SE) for time intervals Δt, and lower **H**, turning angles (mean ± SE); mean angles <90° indicate persistent motion over interval Δt. Lm-RT-m cells move independently from each other, as shown by **I**, angles versus distance of single “steps” co-occurring at the same time (gray points reflect 2% of data to avoid overplotting). The mean angle (blue line) is 90°if bacteria move independently, with deviations indicating crowding effects (if limited to small distances) or global directionality (if systematic).

We next assessed bacterial motility in vitro using video microscopy. Because EGD Lm and Lm^Mu^ were nearly the same in terms of motility (Supp. Fig. S1A,B), we focused on EGD Lm, which has the broadest relevance. Lm-37 and Lm-RT were plated on slides for time-lapse imaging and cell motility was tracked (red dots, Fig. 1C,D, Supp. Movies S1,S2). In contrast to Lm-37, which were predominantly non-motile (<2% motile, Fig. 1C), Lm-RT were highly motile with heterogeneous speeds and long persistent tracks, moving in random directions across the surface of the slide (Fig. 1D,E and Supp. Movies S1,S2). Basic motility parameters for Lm-37 and Lm-RT were calculated and compared (Fig. 1E-I) using *celltrackR* ^29^. Despite large variation in average track speed, Lm-RT motility (∼10 μm/s) dramatically exceeded that of Lm-37 (which were essentially non-motile; <1μm/s Fig. 1E,F). These data confirmed that Lm-RT is highly motile with strongly increased mean squared displacement (motility coefficient M = 19 μm^2^/sec, Supp. Fig. S1C-F) compared to Lm-37 (M = 0 μm^2^/sec). Importantly, Lm-RT maintain this motility even after several hours of subsequent culture at 37°C (Supp. Fig. S1G).

Lm-RT cultures also contained a subset of non-motile or “spinning” bacteria that were stuck to the glass slide. These tracks did not represent motility behavior per se but rather were an artefact of our in vitro system. Therefore, we filtered out these tracks to obtain an idealized “RT-motile” (RT-m) population for further analysis (Supp. Fig. S2). In contrast to the non-motile Lm-37, Lm-RT(-m) tracks were well-described by a persistent random walk model (Supp. Fig. S1) with a motility coefficient M = 19 μm^2^/s in the total Lm-RT population. This motility coefficient increased more than 2.5-fold (M = 52 μm^2^/sec) upon filtering for the motile (RT-m) subset (Supp. Fig. S1A,C), contributing to even larger displacements over time (Fig. 1G). RT-m also had a (slightly) higher persistence time (P = 0.7s vs P = 0.43s or 1.12s vs 0.99s, depending on the analysis method; see Supp. Fig. S1C,D,F). In line with such short-term persistent motion, average turning angles were below 90° for time intervals up to two or three seconds, with lower turning angles observed in the RT-m subset (Fig. 1H). This directional autocorrelation was not due to global directed motion or local alignment due to crowding: average angles between cells were (nearly) equal to 90° (Fig. 1I), suggesting that individual bacteria moved in directions independent of that of other bacteria. In summary, these data show that Listeria cultured at RT display a high, persistent random walk-like motility that lasts for several hours at higher temperatures — suggesting that ingested Lm are predominately motile when they reach the small intestine and invade the epithelium.

### Motility facilitates Lm scanning of the intestinal epithelium and enhances invasion

Next, we examined the bacterial behavior at the epithelial surface using explanted intestinal tissues and 2P microscopy. Sections of B6 mouse intestine were harvested, glued to plastic coverslips, and then sliced open to expose the epithelial surface. Intestines were challenged with murinized Lm-37 or Lm-RT and 2P time-lapse recordings collected to analyze bacterial motility, epithelial adhesion, and invasion (Fig. 2, Movies S3,S4). Non-motile Lm-37 was predominantly found collecting on the top of the mucus layer (10-40 μm above the epithelium) along with red fluorospheres that served as a control to identify regions of the intestine were mucus was thin or absent (Fig. 2A-C). In contrast, motile Lm-RT often penetrated the mucus layer within minutes and multiple bacteria were observed scanning along the epithelium (Fig. 2D-F).

**Figure 2:**
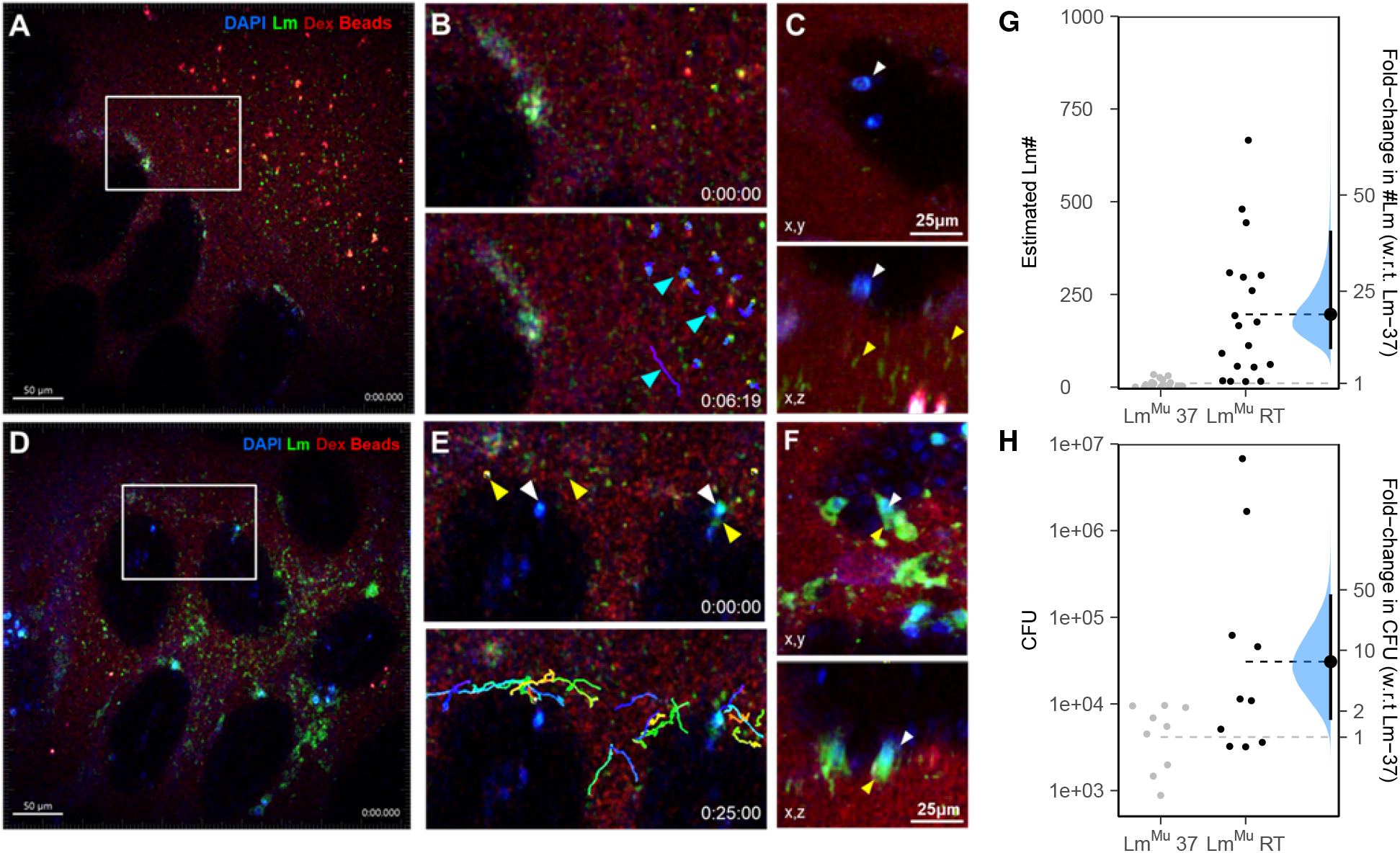
Lm^Mu^ invasion dynamics of mouse intestinal epithelium. **A**, mouse intestine was explanted, challenged with Lm-37 (green) and imaged with time-lapse 2P microscopy. **B**, zoomed views taken from a 25s recording show that Lm-37 are predominantly non-motile (bottom panel, cyan arrows, time encoded tracks) and colocalize with fluorescent beads (pinkish red) trapped at the mucus interface. Rh-dextran (red) was added to confirm the integrity of the epithelium (autofluorescence, blue) and label the fluid phase. **C**, 10min post infection (mpi), Lm-37 (green, yellow arrows) do not accumulate around secreting GCs identified by brightly stained nuclei (white-blue, white arrow). Right and lower panels are orthogonal views projected along the x and y dimensions respectively. **D**, Lm-RT (green, yellow arrows) are motile and penetrate the mucus layer to move along the surface of the epithelium. **E**, zoomed views, within minutes, Lm-RT (yellow arrows) can be seen scanning and adhering near GCs (white arrows). Bottom panel, Lm-RT tracks time encoded (blue to red). **F**, 10 mpi, Lm-RT (green, yellow arrows) accumulates around GCs (bright white-blue nuclei, white arrows). Right and lower panels are orthogonal views projected along the x and y dimensions respectively. **G**, estimated # Lm invading for Lm-37 and Lm-RT was assessed by counting the number of green pixels that overlap with DAPI stained nuclei in the epithelium 10mpi. Invasion was about 19-fold higher (95% CI: 9.8-41) with Lm-RT. Each point represents an image (from 4 mice total). **H**, mice were infected orally with either Lm-37 or Lm-RT and colony forming units (CFU) were determined in the spleen 3 dpi (days post infection). 3 d.p.i., Lm-RT infection yielded 7-fold higher CFU compared to Lm-37 (95% CI: 1.6-45). Each point represents one mouse. In **G**,**H**, horizontal lines represent means for Lm-RT and Lm-37. On the right, plots show the fold-change from Lm-37 to Lm-RT (black dots), along with its bootstrapped distribution (blue) and 95% CI (line segments).

Even in regions where mucus was thin or absent, Lm-37 rarely contacted or attached to the epithelium. However, in intestines challenged with Lm-RT, bacteria accumulated on the epithelium in clumps associated with DAPI stained nuclei, presumably goblet cells. In previous published work, goblet cell nuclei were shown to stain brightly with DAPI during secretion and antigen transcytosis ^30^, suggesting that bacteria associate with goblet cells as previously reported ^12^. We quantified Lm invasion of the epithelium by measuring green pixels (BacLight-Green stained bacteria) that bound to the epithelium over time (Fig. 2G). Few bacteria were found attached to the epithelium initially after challenge. However, within 15min, Lm-RT showed a 19-fold higher attachment compared to Lm-37, which attached infrequently (Fig. 2G, fold-change 95% CI: 9.8-41). We also performed CFU assays on orally infected mice to assess whether motile Lm-RT infects mice more readily than non-motile Lm-37 in vivo. We found that mice infected with Lm-RT had 7-fold higher bacterial burdens in the spleen 3 days after infection than mice challenged with Lm-37 (95% CI: 1.6-45, Fig. 2H). Overall, these results show that Lm-RT, but not Lm-37, actively scan the mouse intestinal epithelium to locate and attach to goblet cells, and that this motility is associated with increased infection.

### Motility promotes Lm interactions with human epithelium and invasion

Mouse infection models provide important in vivo insight into bacterial pathogenesis but, due to the species specificity, standard oral models using Lm^WT^ are inappropriate for studying early host-pathogen interactions in the gut. InlA murinized Lm strains have been used to study oral infection, but murinized Lm has been shown to bind mouse N-cadherin in addition to E-cadherin, potentially affecting invasion specificity ^17^. Therefore, in a complimentary approach, we examined Lm^WT^ invasion in explanted human intestinal tissue biopsies. Human tissue explant systems have the advantage of preserving species-specific invasion mechanisms and host-pathogen interactions and are generally applicable to other intestinal pathogens ^31^.

We used 2P microscopy to assess whether Lm^WT^ motility impacted epithelial invasion in explanted human intestinal tissues. Fresh surgical biopsy specimens were collected from the Washington University DDRCC (Digestive Disease Research Core Center). The muscle layer was removed, and tissue glued with the epithelial surface up on plastic coverslips. Samples were placed in a custom imaging chamber to perfuse the tissue from below with warm oxygenated media. Tissues were infected with either EGD Lm^WT^ grown at 37°C or RT for time-lapse imaging (Fig. 3, Movies S5,S6). The epithelial barrier of explanted human intestine maintained its integrity for several hours as shown by the exclusion of Rh-dextran from beneath the epithelium (Fig. 3A). Human epithelial cells are brightly auto-fluorescent when excited with 800-850nm 2P laser light (Fig. 3A), and goblet cells often appear red due to the uptake of soluble Rh-dextran following mucus secretion, a phenomenon called goblet-cell associated antigen passages ^30^. EGD bacteria co-localized with goblet cells and were found immediately below the epithelium where lamina propria phagocytes reside (Fig. 3B). Human intestine infected with EGD and stained with antibodies to E-cadherin show EGD bound to adherens junctions deep in the epithelium near cells with goblet cell morphology (Fig. 3C).

**Figure 3:**
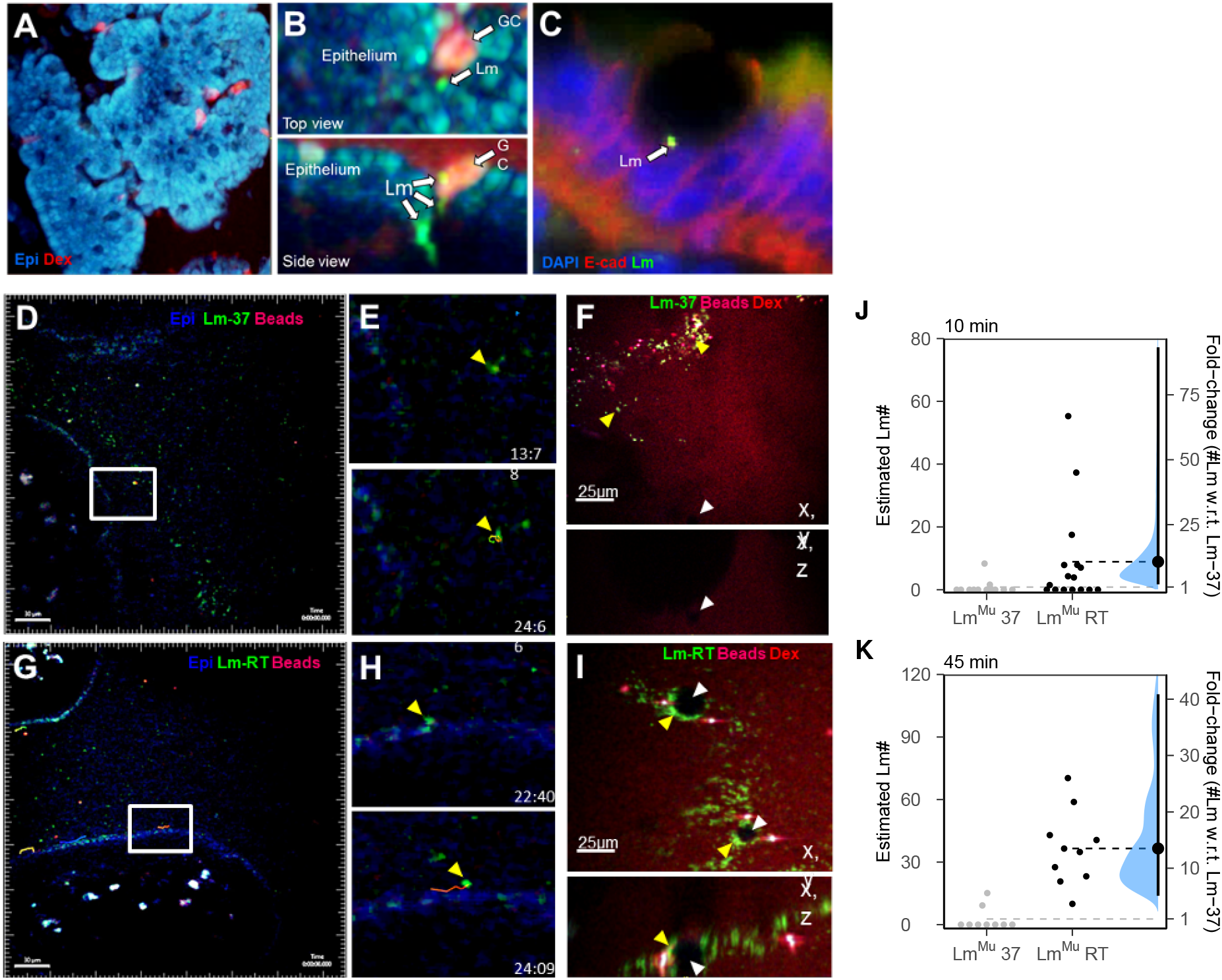
Lm^wt^ interaction dynamics with human intestinal epithelium. **A**,**B** human intestinal resection specimens were challenged with Lm^Wt^ (green) and imaged with two-photon microscopy. **A**, autofluorescence of epithelial cells (blue) excited at 820nm. Several secreting goblet cells have been labeled by Rh-dextran (red) uptake. **B**, Lm-RT (green) can be seen binding to GCs and crossing the epithelium. **C**, Lm-RT (green) binding to E-cadherin (red) near a GC. Epithelial cell nuclei stained with DAPI (blue). **D**,**E**, human intestinal resection specimens were challenged with Lm and imaged with time-lapse two-photon microscopy. **D**, Lm-37 localizes predominately in the mucus layer identified by fluorescent beads (red) above the epithelium (blue, autofluorescence), with **E**, zoomed in views from a time-lapse sequence showing a Lm-37 track (time encoded). **F**, Lm-37 rarely attaches to the epithelium and fails to accumulate around GCs 45mpi. **G**, Lm-RT are often seen penetrating the mucus and scanning along the epithelium, with **H**, zoomed in views from a time-lapse sequence with a Lm-RT track shown (time encoded). Time stamp=min:sec. **I**, Lm-RT (green) attaches to the epithelium and accumulates around goblet cells (dark shadows) forming “rings”. Lm invasion efficiency with Lm-37 and Lm-RT was assessed at **J**, 10mpi and **K**, 45mpi. Invasion was significantly higher for motile Lm-RT (fold-change w.r.t. Lm-37 was 11 at 10 mpi and 14 at 45 mpi, with 95% CIs [2.1-94] and [5.1-41], respectively). Bootstrapped fold-changes displayed as in Fig. 2G,H.

We also infected intestinal explants with EGD-Lm-GFP and analyzed their behavior and interactions with the intestinal mucosa (Fig. 3D-I). Like in the mouse epithelium (Fig. 2), non-motile EGD Lm-37 primarily became trapped in the mucous layer identified by fluorescent beads (Fig. 3F). However, in some areas where the mucus layer was thin or absent, EGD drifted down to the epithelium and attached. In contrast, when intestine samples were challenged with EGD Lm-RT, we observed motile bacteria penetrating the mucus and approaching the epithelium or moving into regions where the mucus was discontinuous (Fig. 3G-I). When motile bacteria encountered the epithelium, they either bounced off and left the field of view or scanned along the epithelial surface (Fig. 3H). We observed several examples of scanning bacteria suddenly arresting and adhering to the epithelium at locations where other bacteria had previously bound. Over 20-30 minutes, bacteria accumulated into green rings in the epithelium (Fig. 3I), presumably as Lm bound to E-cadherin on goblet cells.

We quantified the extent of epithelial invasion by measuring the number of green pixels associated with the epithelium at 10 and 45 minutes after challenge and extrapolating the data to estimate bacterial numbers (Fig. 3J,K, see methods). EGD Lm-RT accumulated significantly more on the epithelium compared to EGD Lm-37 (Fig. 3K). Therefore, the invasion advantage of motile Lm-RT compared to Lm-37 is observed in both the mouse model and the human explant system.

### A layered cellular Potts model shows that bacterial motility can facilitate invasion by driving rapid interactions with target cells

Results in both the human and murine intestinal explant systems suggest that bacterial motility enhances epithelial invasion of “target” cells (i.e. goblet cells). To investigate the consequences of this motility benefit over larger spatiotemporal ranges, we used a computational biology approach to estimate how bacterial motility affects immunological outcomes, such as the number of bacteria that invade target cells or are phagocytosed over time. This allowed us to investigate which host-pathogen interaction parameters determine epithelial invasion efficiency.

We turned to the cellular Potts model (CPM), which can model complex tissues in space and time with realistic cell morphology, motility, and cell-cell interactions ^26–28,32–34^. Our “layered” L-CPM models bacterial motility and invasion dynamics at the epithelium, using simulations with aligned coordinates to simulate the quasi-2D system of bacteria scanning on top of the epithelial surface (Fig. 4A, Movie S7). This approach preserves the relative simplicity of 2D models while also allowing cells to move on top of each other and interact. Importantly, additional CPM layers can easily be added to model more complex interactions, for example adding host phagocytes to simulated host immune response dynamics at the epithelium (see next section).

**Figure 4:**
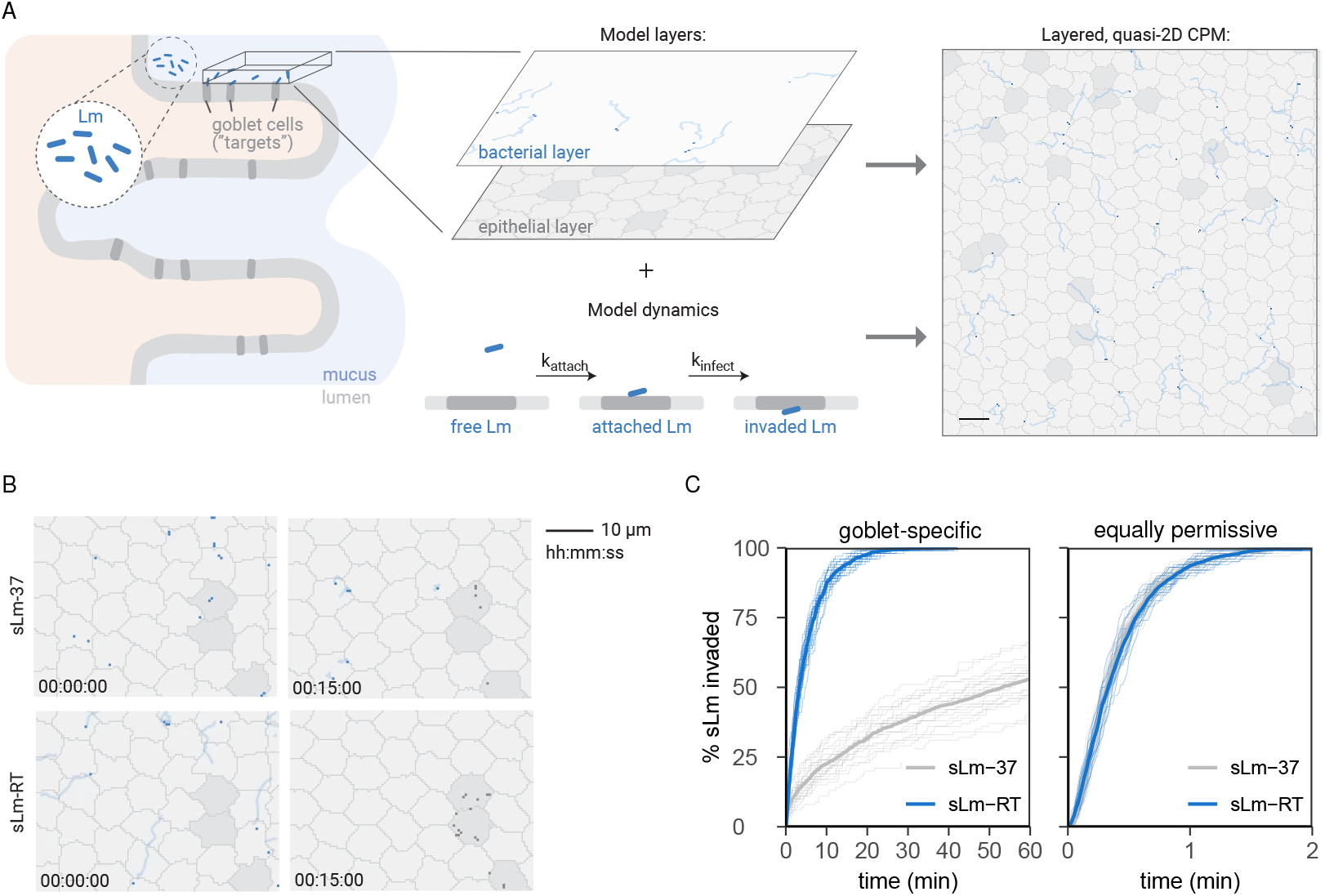
Bacterial motility enhances target cell invasion in a cellular Potts model (CPM) of the epithelium. **A**, The CPM describes the surface of the small intestinal epithelium (left). The model consists of two layers: the gut epithelium (including target cells in dark gray) and a layer above where bacteria (simulated Lm, sLm) move. Bacteria (blue dots) are shown with their traces (light blue). A bacterium that finds itself above a target cell in the epithelial layer can attach to it (rate k_attach_), and subsequently invade (rate k_infect_); both are irreversible processes. These layers together with the invasion dynamics yield the full, quasi-2D model of the gut epithelium (right; scale bar = 10μm). See Methods for details. **B**, Example screenshots of non-motile (sLm-37) and motile bacteria (sLm-RT) at the start of the simulation and after 15 min. **C**, % sLm invading over time, showing 20 individual simulations (thin lines) and average invasion (thick lines) for both non-motile (gray, sLm-37) and motile bacteria (blue, sLm-RT). Left: “goblet-specific” invasion where sLm can only invade goblet “target” cells, right: invasion when all epithelial cells are target cells (equally permissive to sLm).

Our L-CPM is constructed using data obtained from in vitro and in vivo experiments, which give it physiologically relevant spatiotemporal scales as well as quantitative parameters for bacterial motility, invasion efficiency and phagocyte dynamics. First, we measured epithelial cell diameters using in vivo 2P imaging of E-cadherin-CFP (cyan fluorescent protein) mice (gift of Charles Parkos and Ronen Sumagin) and estimated the prevalence of goblet cells (dark goblet shaped cells) in the villus epithelium. These data were used to create the 2D topology of the epithelial layer in the L-CPM (Fig. 4A). In addition, we derived a stereotypical bacterial motility behavior from the in vitro Lm-RT-m motility data (Fig. 1) and chose model parameters to mimic this behavior in a “simulated Lm-RT” (sLm-RT) population (see Supplemental Methods, Supp. Fig. S3), while also simulating a non-motile sLm-37 population. Finally, we viewed time-lapse recordings of motile Lm near goblet cells in the explant intestine models (Movie S3-S6) to estimate a physiological range for the rates of attachment and invasion in the CPM (Fig. 4A, see Supplemental Methods for details).

We then used the CPM to simulate invasion dynamics of sLm-RT as described above (Movie S7). Although Lm-37 were non-motile in vitro, we expect that in vivo, non-motile bacteria would diffuse along the epithelium rather than remain perfectly stationary. Therefore, we modelled sLm-37 with a slight diffusive motion to mimic this behavior and ensure that comparisons with sLm-RT are as conservative as possible (Movie S7). Simulations consistently showed that when sLm have to scan for target cells, sLm-RT motility vastly sped up invasion dynamics compared to the diffusive motion of sLm-37 (Fig. 4B,C). Most sLm-RT invaded the epithelium within 20-30 min (consistent with observations in Fig. 2 and Fig. 3) and in line with published findings (18,20), while in the sLm-37 case, only 50% of bacteria had invaded as late as the 1-hour mark (∼15x later than sLm-RT). The increased invasion efficiency of sLm-RT was a direct consequence of bacterial motility driving interactions with “target” cells (i.e. goblet cells), disappearing in simulations where all epithelial cells were permissive to invasion (Fig. 4C). Indeed, increasing sLm speed facilitated invasion, (Supp. Fig. S4), but once sLm moved fast enough to reach target cells within the 60-minute timeframe, further increasing bacterial speed to “super physiological” levels did not confer any additional invasion advantage.

The finding that bacterial motility confers an invasion benefit is not trivial. While motile bacteria would be expected to find target cells more rapidly, they also, on average, spend less time in contact with the target cells. Therefore, motility could in in theory be *detrimental* to invasion in situations where the rate of target cell attachment is low. However, this detrimental effect did not occur even in simulations with very low attachment rates far outside of the physiological range (Supp. Fig. S5; the 1-hour invasion was consistently about 2-fold higher in sLm-RT than in sLm-37 across all attachment rates). Thus, our model predicts that for physiological bacterial speeds and attachment kinetics, motility helps bacteria locate and invade target cells more efficiently.

### Bacterial motility, not phagocyte motility, drives immunological outcomes in the L-CPM

Next, we examined the impact of host cellular immunity, specifically phagocytosis, on sLm invasion efficiency in the L-CPM. In response to intestinal infection, neutrophils extravasate from submucosal vessels and migrate through the lamina propria towards sites of infection ^35^. Neutrophils can patrol the epithelium directly from the basolateral side as well as undergo transepithelial migration to patrol the apical surface ^36,37^. During the first few hours of infection, neutrophils on the apical surface of the epithelium could lower the initial bacterial burden by phagocytosing bacteria that are scanning or attached to the epithelium, preventing invasion. We tracked and quantified the in vivo motility of LysM-GFP neutrophils in 2D as they crawled along the epithelium in 2P microscopy recordings (Movie S8, Supp. Fig S6,S7). To simulate host pathogen interactions at the epithelium, we expanded our L-CPM by adding a layer of phagocytes with realistic cellular morphology, dynamics and migration behavior based on our in vivo data (Fig. 5A, Supp. Fig. S7, see methods). In this L-CPM, phagocytes can “hunt” sLm as they are scanning the epithelium or immediately after they have attached to target cells, but not yet invaded (see Fig. 4A).

**Figure 5:**
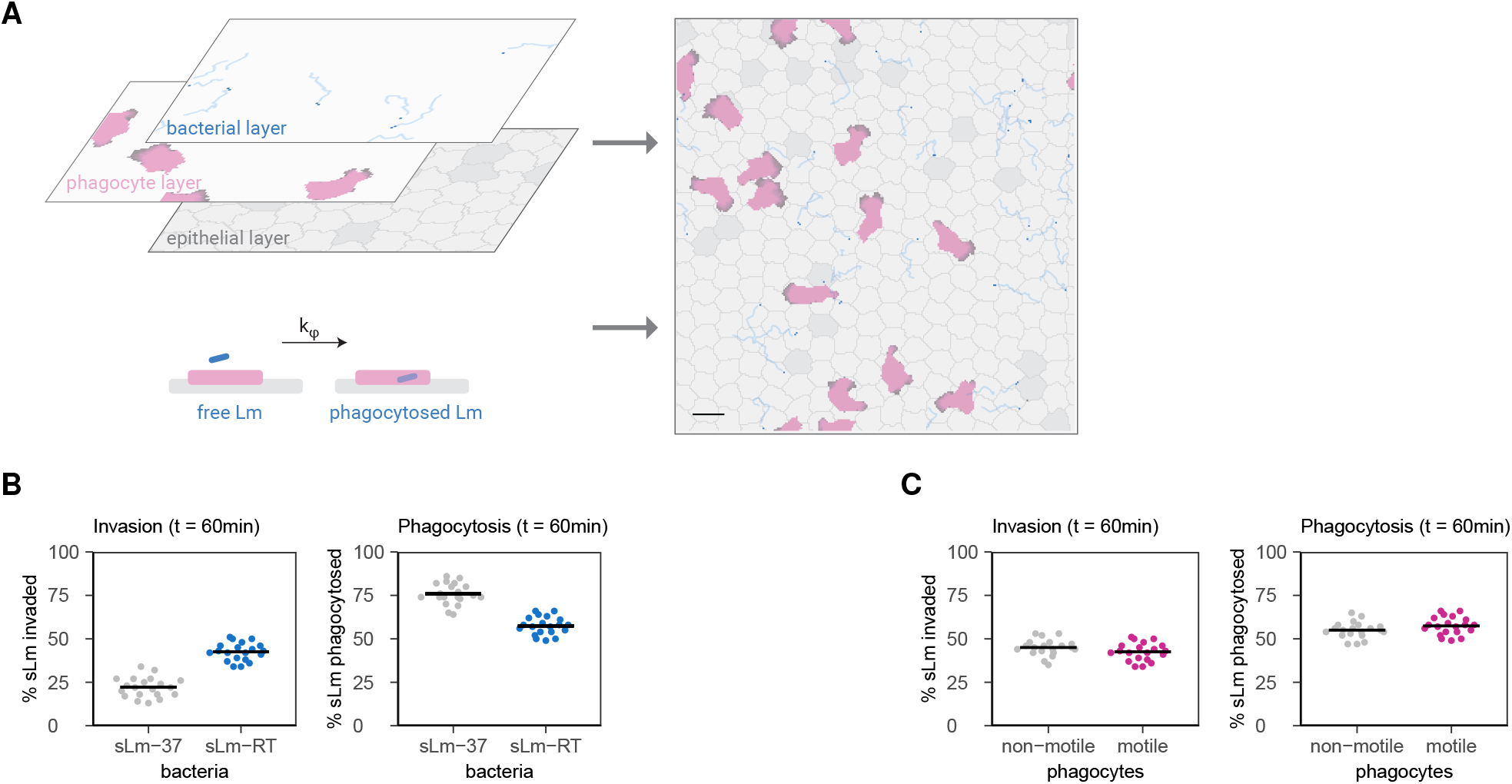
Bacterial motility, but not phagocyte motility, alters immunological outcomes in a CPM model of bacterial scanning the epithelium in presence of phagocytes. **A**, To simulate bacterial phagocyte interactions on the epithelium, the model is extended by adding an extra layer containing migrating phagocytes (left). Phagocytes can phagocytose bacteria that move over them, or that have attached but not yet invaded a target cell. Phagocytosis is irreversible and occurs at rate k_φ_. These layers and dynamics are again combined to obtain the full model (right; scale bar = 10μm). See also Fig. 4 and Supplemental Methods for details. **B**, % sLm invaded (left) and phagocytosed (right) after 60 min for both non-motile (sLm-37) and motile (sLm-RT) bacteria. **C**, the same as panel B, but now comparing motile (pink) versus non-motile (gray) phagocytes, while bacteria are all motile (sLm-RT). Non-motile phagocytes were modelled by setting the parameters λ_act_ = 0, max_act_ = 0 (see Supplemental Methods). For **B**,**C**, horizontal bars indicate the mean of 20 independent simulations.

Although Lm infection induces robust neutrophil recruitment, reliable measures for the number of phagocytes on the epithelium or the phagocytosis rate are currently lacking. Since our focus is understanding how bacterial motility affects host pathogen interactions, we started from a simple, “equal-opportunity” baseline where phagocytes covered roughly the same surface area as target cells and the phagocytosis rate equaled the rate of sLm attachment to target cells (Fig. 5, ensuring that the relative rate differences themselves did not predetermine invasion and phagocytosis outcomes). However, since the true values of these parameters are unknown, we later vary them systematically to assess how they affect our conclusions (see below).

While invasion was again reduced about 2-fold with sLm-37 compared to motile sLm-RT in these baseline simulations, the opposite was true for phagocytosis – which was ∼30% more efficient with non-motile sLm-37 than sLm-RT (Fig. 5B). Non-motile sLm-37 are inefficient at finding (also non-motile) target cells to invade. They would be equally inefficient at finding phagocytes – but since phagocytes are themselves motile, they can still find and sweep up non-motile bacteria from the epithelium to phagocytose (Movie S9). By contrast, phagocytes lose this advantage when faced with the rapidly motile sLm-RT; motile bacteria rapidly encounter both static target cells and slowly moving phagocytes but are also more likely to “escape” from these encounters before invasion or phagocytosis is completed (Movie S9). Thus, motility always helps sLm find other cells faster (be it target cells or phagocytes) – but the success of these encounters depends on whether they last long enough to facilitate the attachment or phagocytosis process.

Because our simple baseline scenario uses “best guesses” for unknowns such as phagocyte numbers and phagocytosis rate, we next examined more thoroughly how sLm fate depends on these parameters. We first varied the prevalence of sLm, target cells, and phagocytes (Supp. Fig. S8, Movies S10-S12). Again, the invasion at 1 hour was consistently about 2-fold higher for motile sLm-RT than for sLm-37 for all phagocyte prevalences (Supp. Fig. S8I) and most challenge doses (Supp. Fig. S8A). The magnitude of this motility-dependent invasion benefit only varied with the number of target cells (Supp. Fig. S8E) – consistent with the hypothesis that motility helps Lm find target cells when these are rare.

While challenging the epithelium with more sLm mostly did not alter the fraction of sLm invaded or phagocytosed, additional target cells or phagocytes shifted Lm fate towards more invasion or phagocytosis, respectively (Supp. Fig. S8, Movies S10-S12). Indeed, when sLm can either invade or be phagocytosed, target cells and phagocytes essentially compete for bacteria; for motile bacteria, it is then the relative surface area of target cells and phagocytes that determines which type of interaction partner it most likely encounters first (Supp. Fig. S8H,D,L). Increasing the rates of target attachment or phagocytosis similarly shifted the invasion-phagocytosis balance – and in contrast to cell prevalences, the choice of these rates *did* affect the strength of the motility-dependent invasion benefit of sLm-RT (Supp. Fig. S9). Still, sLm-RT retained its invasion benefit over sLm-37 in most cases, as this trend was reversed only in cases where phagocytosis was >3x faster than attachment. Thus, sLm motility drives both target cell and phagocyte encounters and almost always confers an invasion benefit, but both this benefit and the invasion-phagocytosis balance depend on quantitative parameters (such as numbers of target cells and phagocytes present, and the relative rates of attachment and phagocytosis). Nevertheless, we conclude that motile bacteria retain their invasion benefit over non-motile bacteria in the presence of phagocytes – unless phagocytosis is over three times as fast as bacterial attachment and invasion.

Finally, a general but striking conclusion followed from this model: bacteria move so much faster than phagocytes that phagocytes are fundamentally incapable of “hunting” motile bacteria – whether that is through random migration or chemotaxis. Rather, phagocytes rely on bacterial motility to drive capture and phagocytosis (Movie S9). Indeed, while sLm motility strongly affected invasion and phagocytosis (Fig. 5B, Supp. Fig. S10A-C), *phagocyte* motility did not, except when sLm were also non-motile (sLm-37, Fig. 5C, Supp. Fig. S10D-F). Thus, our model predicts that once phagocytes have reached the epithelium, their motility will have negligible impact on immunological outcomes during infection with motile bacterial pathogens.

## Discussion

### Lm are likely motile when they reach the gut epithelium

Many important human bacterial pathogens are highly motile (Lm, *E. coli, P. aeruginosa*) ^24,25^, yet few studies have assessed the impact of bacterial motility on infection ^24,25,38^. Our study suggests that Lm motility enhances epithelial infection by allowing bacteria to efficiently scan the epithelium for target cells that are permissive to infection. This hypothesis is based on the observation that Lm preferentially invades the host via goblet cell transcytosis ^12^ and that human food-borne infections typically result from Lm living at low temperatures (≤RT), which are flagellated and motile. Indeed, we confirmed that Lm-RT are flagellated and highly motile and showed that their motility resembles a random walk with strong directional persistence.

Previous studies have examined the role of Lm motility in infection, but the findings have been inconsistent. Way et al. found that bacterial burdens and immune responses were similar between Lm flagellar null mutants and WT bacteria in both i.v. and oral gavage infection mouse models ^39^. In other work, however, O’Neil et al. reported that infection with flagellated Lm outcompeted non-flagellated bacteria 6-16hrs after oral challenge ^22^. Experimental systems could account for these different findings, since mice infected by i.v. injection bypass the epithelial invasion step, and infection by gavage can produce small tears in the esophagus and lead to direct entry of Lm into the circulation in contrast to oral ingestion approaches ^15,40^. Furthermore, most Lm strains isolated from patients are indeed non-flagellated, whereas human infections result from eating contaminated foods (stored at less than 37°C), yielding bacteria that are presumably flagellated and motile when ingested. One potential caveat is that Lm could rapidly downregulate flagella expression after ingestion and would therefore enter the small intestine in a non-flagellated state. Indeed, flagellin is a potent TLR ligand, and lack of expression by Lm-37 could confer an infection advantage by making Lm less immunogenic. However, we found that Lm-RT (both EDG and 10403s strains) retain motility for several hours after incubation at 37°C (79% motile at 2h and 56% at 3h, Supp. Fig. S1E), suggesting that Lm likely remain motile upon entering the small intestine.

### Bacterial motility facilitates invasion

2P imaging of explant tissues revealed that when motile Lm reached the gut epithelium, their motility enhanced bacterial penetration of the mucous layer above villi and allowed bacteria to explore the surface of the epithelium. Moreover, motility allowed pathogenic bacteria to enter sites where the mucus barrier is imperfect, and thus to gain access to the epithelium.

As informative as tissue explant systems are, they lack immune cell recruitment, blood flow and enervation, and do not provide downstream immunological outcome measures. In complementary computer simulations, we therefore integrated quantitative measurements of cell sizes and motility obtained from imaging experiments, and assessed invasion outcomes over a wide range of infection scenarios (e.g. bacterial challenge doses and motility parameters) and time scales that are not feasible to image directly. Consistent with our two-photon imaging results, CPM simulations predict that bacterial motility enhances invasion by allowing bacteria to scan the epithelial surface and quickly locate preferred target cells for invasion. This invasion advantage associated with bacterial motility holds in most, but not all conditions tested – for example, it disappears when cell specificity is irrelevant (i.e., all epithelial cells are equally permissive for invasion), or when phagocytosis is so efficient that motile bacteria are phagocytosed before they can invade (very high rates of phagocytosis). Nevertheless, our model predicts that this effect is surprisingly robust: in absence of phagocytes, invasion at 1 hour was consistently ∼2-fold higher for motile bacteria, regardless of the rate by which they attach to target cells. Upon arrival of phagocytes, the size and direction of the effect does depend on attachment and phagocytosis rates – but it did persist at least qualitatively in almost all scenarios. Thus, unless there is reason to believe that phagocytosis rates are at least 3-fold higher than the rate of Lm attachment to target cells, our model predicts that motile bacteria have an invasion advantage once they reach the epithelium.

### Bacterial motility, not phagocyte motility, drives phagocytosis

Aside from showing how bacterial motility facilitates invasion, our quantitative results also challenge the fundamental paradigm of phagocytosis as “predators”. By simulating bacterial and phagocyte dynamics to scale, our CPM clearly shows that neutrophils cannot possibly “hunt” motile bacteria – which move at ∼120X their own speed. Surprisingly, bacterial motility *can* drive an increase in phagocytosis in our CPM – at least in conditions where phagocytosis is more efficient than invasion. This suggests that rather than pursuing their prey, phagocytes may act more like “fly paper”, capturing and ingesting motile bacteria in the local environment through random collisions. Indeed, goblet cells and phagocytes can be seen as as two competing traps, whose relative “surface area” (abundance of goblet cells/phagocytes) and “stickiness” (rates of goblet attachment/phagocytosis) determine the balance between attachment/invasion and phagocytosis.

We note that even though phagocyte motility had a negligible role in capturing motile bacteria in our simulations, this does not suggest that motility is entirely dispensable. Phagocyte motility is clearly required for many aspects of phagocyte function including extravasation from blood vessels and trafficking to sites of infection. Furthermore, our simulations showed that non-motile bacteria *can* be “hunted” actively by phagocytes. This scenario would apply to skin infections with S. aureus or S. pyogenes ^41,42^ (both non-motile cocci), where phagocytes are important for controlling bacteria growth and dissemination through phagocytosis and cytokine secretion.

### Future directions

Our L-CPM allows us to investigate a wide range of infection scenarios by directly controlling the number of phagocytes and bacteria in the simulations over time. However, like any model, it has its limitations. First, it is worth noting that the 2D interactions modelled by our L-CPM are (due to spatial constraints) more likely to drive random phagocyte-bacteria collisions than 3D interactions would. Although this is not a problem in the specific case of the epithelium modelled here (which essentially *is* a 2D system), care must be taken when translating these findings to other tissues. Still, this 2D topology is relevant to many barrier surfaces such as the upper and lower airways, the skin, and the eyes, as well as the lining of sinuses in the liver and spleen.

Second, our simple model does not account for host responses below the epithelium (e.g., lamina propria macrophages) or the effect of the mucus layer containing normal flora above the epithelium. These are important phenomena that can be investigated in future iterations of the model with additional 2D CPM layers. Indeed, this is a significant strength of our layered CPM: it can be readily modified to accommodate more complex host-pathogen systems, without the complications arising from rendering every component of the system in 3D.

One such model extension could be aimed at examining the effect of bacterial motility over longer time scales. We here focused the dynamics of bacterial invasion and phagocytosis during the first hour after infection, but a slightly more complex model could extend this period to multiple hours. Such an extended model should take into account that in vivo, bacteria take several hours to transit down the intestine after oral infection, and that phagocytes are recruited in multiple steps; neutrophils extravasate from submucosal vessels and migrate through the lamina propria towards sites of bacterial infection, where they can patrol the basolateral side of the epithelium or undergo transepithelial migration to patrol the apical surface ^36,37^. The L-CPM could implement these dynamics by gradually increasing bacteria and phagocyte numbers over time. The current model suggests that the earliest Lm invasion events occur too rapidly for neutrophils to be recruited – but by simulating neutrophil recruitment kinetics, the model could test under which conditions recruitment farther down the digestive tract (ahead of infection) could reduce Lm invasion and bacterial burden during the later stages of infection. Furthermore, we often observe blood in the lumen of Lm infected mouse intestines. Since neutrophils are abundant in blood (300-500 polymorphonuclear neutrophils per μl of blood in C57/B6 mice ^43^), an interesting future avenue would be to study the role of bleeding in anti-bacterial host responses.

For future work, it will also be important to develop in vitro and/or in vivo systems in which the rates of phagocytosis, epithelial attachment and invasion can be measured more precisely than we do here. With more robust estimates of phagocytosis, attachment, and invasion rates, future CPM simulations could also explore the effect of bacterial opsonization by antibody or complement on phagocytosis efficiency, bacterial motility and invasion efficiency.

## Conclusion

In summary, our CPM simulations highlight the competition between phagocytosis and invasion in the system and suggest that this interaction could be crucial to infection outcomes. Throughout a wide range of tested parameters, our model supports the idea that Lm motility confers an invasion advantage because it allows Lm to quickly locate its preferred target cells (i.e. goblet cells). We believe that our L-CPM-based modeling approach is highly accessible and adaptable and thus could be applied to study cell dynamics in a wide range of systems, including cellular immunity to pulmonary or skin infections, host microbiota interactions, bacterial biofilm formation, antigen presentation, tumor immunology and neuroimmune cross talk.

## Methods

### Mouse strains

Mice were bred in house or purchased from JAX labs. B6 mice were bred and housed under specific pathogen–free conditions in the animal facility at the Washington University Medical Center. The use of all laboratory animals was approved and performed in accordance with the Washington University Division of Comparative Medicine guidelines.

### Bacterial culture

Lm EGD ^44^, PNF8-GFP ^45^ and Lm^Mu^ strains ^14^ were stored as frozen glycerol stocks (∼1×10^9^/ml) at -80°C. Bacteria were cultured in BHI medium and harvested during log phase growth for inoculation of tissue and mice ^46^. Lm concentrations in culture were estimated from standard growth curves by measuring optical density at 600nm.

### Colony forming unit (CFU) assays

No experiments used death as an endpoint and the Lm challenge doses used are well tolerated in B6 background mice. Mice were given P2R inhibitor and PBS vehicle controls then infected with 2×10^7^ Lm^Mu^ by direct luminal injection. Mice were sacrificed 4h.p.i. and the ileum, mLM and spleens harvested and incubated in CO_2_-independent media containing 25 μg/ml gentamicin for an hour ^47^. Tissues were transferred to a RINO tube containing 0.6ml DPBS and 5 SSB32 stainless steel beads and homogenized, diluted and plated onto BHI agar plates and ChromAgar Listeria selective plates. 1-2 days after plating, colonies were counted and CFU analyzed in R (see “Statistics and group sizes in explant experiments” below). For CFU assays, human and mouse tissues were incubated with gentamicin in DMEM to kill extracellular Lm. Tissues were washed 3x, homogenized and serial dilutions plated on Lm selective media to calculate CFU ^47^. In some experiments, the intestine was fractionated into epithelial and LP populations by incubation with EDTA/gentle shaking and cells isolated by flow sorting as previously described ^30^ to allow CFU measurements for cell subsets.

### In vitro bacterial motility analysis

Two 2ml culture of Lm-InlA was grown in 14ml polystyrene round-bottom tube (Falcon Cat No 352057) at RT and 37°C with shaking at 200rpm to OD600 around 1.2. The sample was diluted 1:10 with BHI and 8 μl of 1:10 dilution was used on a non-charged slide (Globe Scientific Cat No 1324W) to check motility. A 2D time lapse video was recorded for 15 seconds with 250ms time interval (60 time points) and 100ms exposure using an Olympus IX51 inverted microscope with 20X objective and phase dichroic filter. The 2D time lapse video was converted to an Imaris file using Imaris 9.5 (RRID:RRID:SCR 007370) and cells were tracked, and percent motile cells were calculated using a 4 μm track displacement length filter. Tracked cells were imported into *celltrackR* ^29^ (RRID:SCR 021021), after which mean track speed, mean squared displacement (MSD), and autocovariance/turning angles were computed as described in the package documentation ^29^ and compared between RT and 37°C. As a measure of uncertainty in the difference in population mean speeds, the following was repeated 10000 times: individual track speeds from both datasets were pooled and sampled with replacement to obtain a resample of equal size. Within that sample, the difference in population means was then determined to obtain the bootstrap distribution of 10000 estimates and its 99% confidence interval (Fig. 1F).

Lm-RT were then filtered for “motile” tracks, Lm-RTm, and their mean squared displacements were fitted to obtain the motility coefficient M and the persistence time P (see Supplemental methods for details).

### Electron microscopy of bacterial flagella

Murinized Listeria Monocytogenes (Lm-InlA) were grown in BHI media at RT and 37°C overnight with shaking at 200rpm. OD600 was measured and overnight culture was diluted to OD600 = 0.6 with BHI. Motility was confirmed by taking a 2D time lapse video recorded for 15 seconds with 250ms time interval (30 time points) and 100ms exposure using an Olympus IX51 inverted microscope with 20X objective and phase dichroic filter. Lm-InlA grown at RT was motile while Lm-InlA grown at 37°C was non motile. OD600=0.6 culture bacteria were fixed with 1% glutaraldehyde (Ted Pella Inc., Redding CA) and allowed to absorb onto freshly glow discharged formvar/carbon-coated copper grids for 10 min. Grids were then washed in dH2O and stained with 1% aqueous uranyl acetate (Ted Pella Inc.) for 1 min. Excess liquid was gently wicked off and grids were allowed to air dry. Samples were viewed on a JEOL 1200EX transmission electron microscope (JEOL USA, Peabody, MA) equipped with an AMT 8-megapixel digital camera (Advanced Microscopy Techniques, Woburn, MA).

### 2P imaging of neutrophil recruitment in vivo

LysM-GFP reporter mice were anesthetized, and an incision made in the lower abdomen to expose the ileum for intraluminal infection, which better synchronizes Lm invasion events for 2P imaging experiments. Mice were placed in a warmed imaging chamber and given fluids for experiments lasting more than 2hr. A region of the ileum containing 1-2 Peyer’s patches was secured to a plastic coverslip for support ^30^. Blood vessels were labeled by r.o. injection of 20 μl of 655nm non-targeted QD in 40 μl of PBS. 2P imaging was performed with a custom-built dual-laser 2P microscope ^48^ equipped with a 1.0 NA 20x water dipping objective (Olympus). Samples were excited with a Chameleon Vision II Ti:Sapphire laser (Coherent) tuned from 750-980nm depending on the experiment and fluorescence emission detected by PMTs simultaneously using appropriate emission filters to separate SHG and the various fluorescence signals.

Video-rate 2P imaging (30f/sec) was performed 50-200 μm beneath the serosal surface of the ileum. Video-rate imaging was used to assess neutrophil (LysM-GFP) recruitment from 0-2 hpi at multiple trafficking stages and 3D time-lapse imaging (220 × 240 × 50μm, every 20-30 sec) for to record the interstitial of leukocytes to the epithelium. Multi-dimensional data sets were rendered, and cells tracked using Imaris (Bitplane).

### 2P imaging of bacterial epithelial scanning, attachment, and invasion in explanted mouse intestine

Mice were euthanized and small intestine harvested and placed in CO_2_ independent media. Intestines were glued to plastic coverslips using VetBond adhesive and gently cut open longitudinally to expose the luminal surface of the epithelium. Explanted tissues were placed in a custom imaging chamber and covered in DMEM without phenol red containing 2μm red fluorospheres (em 625nm) to identify the mucous layer and tetramethyl rhodamine (TMR)-Dextran to assess epithelial integrity and goblet cell secretion. In some experiments, E-cadherin-CFP mice were imaged to assess epithelial cell numbers and dimensions in the intestinal villi ^35^. Tissue explants were infected with 1×10^7^ Lm and imaged with 2P microscopy to record Lm scanning behavior as wells as epithelial attachment and invasion. Lm attachment was assessed by examining the epithelium for the presence of Lm and then quantifying attachment by measuring the number of green pixels and applying a conversion factor of 23.6 voxels/Lm (3D).

### 2P imaging of bacterial epithelial scanning, attachment in human intestine

Fresh human small intestine resection specimens (∼4-9 mm^2^,∼8 samples total) were obtained through the WUSM Digestive Diseases Research Core Center and placed in CO_2_ independent media at 4°C for transport to the imaging lab. The muscularis was trimmed away using surgical scissors to prevent intestinal contraction artefacts and the tissue glued to plastic cover slips using VetBond adhesive (3M), soaked in 10kD TMR-dextran and placed in a custom imaging chamber epithelial surface facing up. Epithelial integrity was assessed by verifying that TMR-dextran is excluded from the LP and that epithelial layer continuity is preserved. The epithelium was also assessed using 800nm excitation, which induces a strong intrinsic fluorescence signal (<480nm) in epithelial cells. 1×10^7^CFSE-labeled EGD or PNF8-GFP Lm or 1.0μm green/yellow carboxylate-Fluorspheres (ThermoFisher) in 25 μl of DMEM was pipetted on epithelium. The tissue was placed in warm oxygenated DMEM maintained at 37°C under slow flow (<1ml/min) for 2P time-lapse imaging. Tissues remained viable for 2-4 hours. Lm attachment was assessed by examining the epithelium for the presence of Lm and then quantifying attachment by measuring the number of green pixels and applying a conversion factor of 9.92pixels/Lm.

### Immunofluorescence microscopy (IFM) of tissue sections

After 2P imaging, tissue sections were fixed and analyzed using epifluorescence microscopy to enumerate neutrophils and monocytes in the tissue. The ileum was harvested 4 hpi., fixed in 4% PFA overnight, embedded in OCT compound (Sakura Finetek) and 5-15 μm cryostat sections cut. Sections were stained with antibodies to E-cadherin and DAPI. All antibodies and isotype matched control antibodies are commercially available (BioLegend, Invitrogen).

### Statistics and group sizes in explant experiments

In the explant studies shown in Figs. 2 and 3, group sizes consist of 3-5 mice or human tissue explants. Measurements were made in a blinded fashion whenever possible. Results were reproduced in a minimum of two independent experiments. Rather than computing p-values, in accordance with more recent recommendations ^49^ we instead report effect sizes along with their 95% confidence interval (CI) as estimated via bootstrapping. To compare the #Lm attached to goblet cells, we used the bootstrapped fold-change of the means (Lm-RT/Lm-37), which we considered the most interpretable and meaningful effect size estimate. Since the distribution of CFU values (Fig. 2H) was extremely skewed, data were plotted on a logarithmic axis and fold-changes were computed on this scale as well (i.e., as the exponent of the difference in means of log-transformed values). Bootstrapping was performed in R (v4.1.3) using R packages *boot* (v1.3.28) and *simpleboot* (v1.1.7), with 10^5^ bootstrap samples and using the percentile-based CI.

### Cellular Potts model of the epithelium

Our cellular Potts model (CPM) of the epithelium (Fig. 4,5) was built in Artistoo ^50^. An interactive web version is available at: https://ingewortel.github.io/2022-listeria-goblets. This web tool requires no prior knowledge or installation and allows readers to explore the model with different settings and export results. All code required to reproduce the modelling results in this manuscript will also be made available on Github upon publication of this manuscript.

The model simulates a 125×125μm patch of epithelium with periodic borders, such that cells crossing the border on the right will re-enter the field on the left and vice versa. Separate layers describe the dynamics of the phagocytes, the epithelium, and the Lm scanning it (Fig. 4,5). Each layer contains its own CPM, essentially an image consisting of different “pixels”, reflecting bits of space that either contain a specific cell or only empty background. We chose a spatial resolution of 2 pixels/μm; at this scale, Lm typically occupy 1-2 pixels whereas epithelial cells and phagocytes are resolved in more detail (further increasing this resolution would only slow down the simulation unnecessarily). The model then changes over time in simulated “monte carlo steps” (MCS, here 1MCS = 1 sec).

Every MCS, cells first move within their own respective layers, modelled to match the motility of real cells (neutrophils/bacteria; based on published models ^27,28,51^ respectively; see “CPM dynamics”, “Bacterial CPM”, and “Phagocyte CPM” in the Supplemental methods for details). To account for the much higher motility of bacteria, the bacterial CPM was run for v_rel_ =150 steps/s. After migrating *within* their respective layers, cells then interact *between* the layers, letting bacteria either: (1) be phagocytosed by overlapping phagocytes (rate k_φ_), (2) attach to overlapping goblet cells (rate k_attach_), or (3) fully “invade” the goblet cells they are attached to (rate k_infect_). For further details on these processes, we refer to the Supplemental methods (“invasion and phagocytosis dynamics”). For an overview of model parameters and an explanation of how they were selected, please refer to the Supplemental methods and Table S1 in the Supplemental materials.

### Simulation analysis

All simulations were performed for a total duration of 1 hour (=3600 MCS in the epithelial/phagocyte models, and 3600*v_rel_ steps in the bacterial model). Events of invasion and phagocytosis were tracked over time and further processed to obtain outcomes (%bacteria invaded/phagocytosed and %target cells infected). These outcomes were plotted (using R) as mean±standard deviation (SD) or standard error (SE) of 20 independent simulations. This number was sufficient that the SE was typically not or barely visible underneath the plotted lines, reducing uncertainty to a negligible level.

## Supporting information

overview of supplemental movies

## Supplemental methods

Supplemental methods containing full details about the model construction and the selection of model parameters are available in the Supplemental material.

## Author Contributions

IW and MJM designed the research and the computational model. SK performed and analyzed CFU assay, electron microscopy of bacterial flagella, and in vitro bacterial motility assays. Two-photon imaging experiments and analysis were performed by MJM, SK and EI. IW implemented the L-CPM model using the parameters from 2P imaging data to simulate host-pathogen dynamics. AYL assisted with testing and analyzing simulations. IW and MJM wrote the manuscript.

## Acknowledgements

We would like to thank Johannes Textor for many helpful discussions and his expert advice. We would like to thank Charles Parkos, Ronen Sumagin, Vero Azcutia Criado and Mathias Kelm for the E-cadherin-CFP mice. We would like to credit the WashU In Vivo Imaging Core for 2P microscopy, Wandy L. Beatty and the Molecular Microbiology Imaging Facility for electron microscopy, and Lihua Yang for technical help with immunofluorescence microscopy. Finally, we thank Jérémy Postat for helpful comments.

## Supplemental Materials

### Supplemental Methods

#### Filtering for motile Lm

To filter Lm-RT “motile” tracks (Lm-RTm), each track was modelled either as (i) a single gaussian distribution of cell positions (ignoring the links between coordinates, with mean the average position and SD σ=1μm), or (ii) two gaussians of unlinked positions (one for the first *m* coordinates and one for the rest). The bayesian information criterion (BIC) was then computed for model (i) versus the possible models (ii) for all choices of *m*, and compared using ΔBIC=BIC_i_-BIC_ii_. The more positive this number gets, the more evidence there is for model ii over model i – i.e., the more evidence that the cell has moved. We then selected tracks with ΔBIC>50, an empirically determined threshold that separated clearly motile versus clearly non-motile tracks reasonably well (Supp. Fig. S1).

#### Estimating motility statistics

To estimate the motility coefficient *M* and the persistence time *P* of the Lm-37 and Lm-RT(-m) populations (Supp. Fig. S2), we computed mean squared displacements using *celltrackR* ^29^ (RRID:SCR 021021), focusing on the initial part where Δt<5sec (where MSD values are still based on many independent tracks and not biased by artefacts from cells leaving the imaging window ^52^. These data were then fitted using the (2-dimensional) Fü rth’s equation to obtain *M* and *P*:

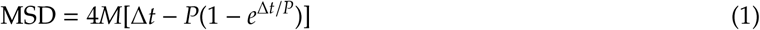

Data were fitted in R using the function nls (port algorithm, starting values [D=10, P=0.005], lower limits [D=0, P=0.001]). This process was bootstrapped to obtain a measure of uncertainty for the estimates of *M* and *P*: each of 1000 bootstrap rounds, tracks were sampled with replacement to get a new track dataset of the same size as the original, and the process above was repeated to obtain the distributions of estimated *M* and *P*.

The persistence *P* was also estimated from the autocovariance, using a similar procedure: autocovariances were computed in *celltrackR*, again only for the initial Δt<5sec, and fitted to an exponential decay function:

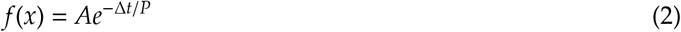

again using nls (port algorithm, starting values [P = 0.5, A = half of the autocovariance at Δt=1], lower bounds [A = 0.001, P = 0.01]).

#### CPM dynamics

For a full description of the CPM we refer the reader elsewhere ^32–34^, but we briefly describe the main dynamics here. At any point in time, CPM pixels *p* can only belong to one cell or background, but these pixel identities σ_*p*_ can change over time as follows: a randomly selected “source” pixel *p*_*s*_ tries to copy its identity into that of a (randomly chosen) neighboring “target” pixel *p*_*t*_. If this succeeds, *p*_*t*_ becomes part of the cell of *p*_*s*_ (changing σ_(_*p*_*t*_) → σ_(_*p*_*s*_)); otherwise nothing changes. The success rate of these copy attempts depends on their effect on the “Hamiltonian” *H*, an energy function defined by the modeler (see below). If the proposed change is energetically favorable (Δ*H* <0), it always succeeds (*P*_copy_ = 1), otherwise, its success probability becomes:

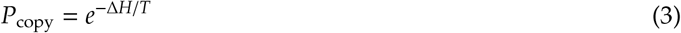

Every MCS, this process is repeated once for every pixel in the model. The temperature *T* determines how much noise (unfavorable changes) the model permits. The model behavior is thus mostly governed by the choice of (Δ)*H*. Below, we list the most important properties and parameters for each model layer.

#### Epithelial CPM

The epithelium was modelled using a standard equation for Δ*H*, yielding a basic CPM ^26^ with 5 parameters. The cell volume *V* (an area in our 2D model), was chosen to match the typical interface of epithelial cells with the mucus/lumen. Based on images of epithelial cells (Supp. Fig. S6A), we can roughly model these as circles with a radius of 8.4μm, yielding an area of ∼55.4μm^2^ = (2pix/μm x 55.4μm) x (2pix/μm x 55.4μm) ≈ 222 pixels. The other parameters determine epithelial dynamics: *J*_cell,background_ = 20, *J*_cell,cell_ = 30, λ_volume_ = 50, *T* = 20.

Target cell frequency in the model was estimated based on our own observations and the published work of others ^53,54^ that showed goblet cells represent between 4-12% of epithelial cells in the mouse small intestine. Varying target cell percentage within this range has only modest effects on model outcomes as the epithelium is relatively static compared to the phagocytes/bacteria.

Using these parameters, 13 × 13 = 289 epithelial cells were seeded, equally spaced, on the CPM. Cells were initialized as circles with a 6-pixel radius and, before the start of the simulation, allowed to equilibrate for 20 MCS to expand and cover the entire grid. *N*_goblet_ cells were then randomly appointed target cells representing “goblet cells” that can potentially be invaded by bacteria unless otherwise mentioned, we used *N*_goblet_ = 20 so that 7% of the epithelial surface is occupied by goblet cells, which is within range of experimental values ^53^.

#### Bacterial CPM

In the bacterial CPM, sLm were modelled as cells that must always occupy either one or two (neighboring) pixels (2-4μm); this roughly matches Lm size. By sequentially adding a new neighboring pixel to the cell and retracting the old pixel, these bacteria can move; but without further constraints this motion is diffusive only. To mimic the directional persistence observed in real bacteria, we used a slight modification of the Beltman et al. model ^51^:

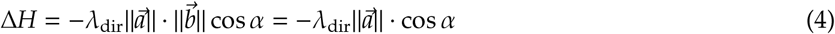

Here, 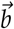 is the normalized displacement vector of the bacterium over the last Δt steps (because of the normalization, its length 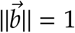. 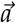 is the vector pointing from the source pixel *p*_*s*_ to the target *p*_*t*_; unlike in Beltman et al. ^51^ it is not normalized, reflecting the slightly larger distance of copy attempts over the diagonal compared to copy attempts in vertical/horizontal directions 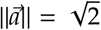 pixels in the former case and 1 pixel in the latter). αis the angle between 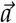 and 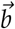, its cosine is +1 when the copy attempt perfectly aligns with the previous cell direction, and -1 when the two are polar opposites. Thus, copy attempts in the direction of previous movement are favored with a negative Δ*H* whose magnitude depends on the parameter λ_dir_.

This model yields motility that roughly matches the persistent random walks exhibited by motile bacteria. The most important parameters controlling this motion are Δt (controlling the persistence time over which the previous cell direction is taken into account) and λ_dir_ (containing the strength of this directionality). Because Lm move at much higher speeds than phagocytes do, the temporal resolution of the bacterial CPM was set higher than that of the rest of the model: for every MCS of the main model, *v*_rel_ steps of the bacterial model are performed. Thus, *v*_rel_ is another model parameter scaling bacterial speed relative to other processes. Δt, λ_dir_, and *v*_rel_ were fitted to match in vitro Lm motility; see “Lm motility parameters” below.

At the beginning of each simulation, *N*_bac_ bacteria were randomly placed in the field with random initial directions. Except in simulations where challenge dose was varied, we used *N*_bac_ = 100.

#### Lm motility parameters

To model Lm-RT motility,λ_dir_, Δt and *v*_rel_ were chosen to match motility statistics of in vitro Lm-RT-m motility data. Because some motility measurements are influenced by imaging window size ^52^, a separate bacterial layer simulation was set up to match the imaging window of the Lm motility data (348.5 × 261.4μm = 697 × 523 pixels), with periodic borders. 50 bacteria were simulated for 30 x *v*_rel_ steps (=30sec) at a given set of parameters, recording centroids every 10 steps. Simulated tracks were then (1) split whenever cells crossed the periodic boundary, to mimic cells entering and leaving the field of view, and (2) interpolated to the same framerate as the original data (every 0.25sec). *T* was (arbitrarily) fixed to 20. λ_dir_, Δt and *v*_rel_ were tuned manually until simulated cells had similar speed distributions, MSD, and autocovariance as vitro Lm-RT (computed in *celltrackR* ^29^, RRID:SCR 021021, Supp. Fig. S3). This yielded λ_dir_ = 40, *v*_rel_ = 150 steps/sec, and Δt=60 steps (on the bacterial CPM, which is 60/*v*_rel_ = 60/150 = 0.4sec in real time). To model non-motile Lm-37, λ_dir_ was set to 0 (ensuring that Δ*H*=0 regardless of Δt; removing all directional persistence). *v*_rel_ was set to 1 to allow Lm-37 some slight diffusive motion, but nowhere near as fast as Lm-RT.

#### Phagocyte CPM

Phagocytes were modelled using the Act-CPM, which has been shown to capture realistic mammalian cell shapes and motility in the CPM ^27,28^. For details we refer to these publications. The cell volume *V* was set to 314 pixels, corresponding to the 78.5μm^2^ area obtained for cells with a diameter of ∼10μm (roughly the scale of the cells in the neutrophil imaging data). The perimeter *P* was set to 230 to get a reasonable circumference for that cell size. Most other parameters were chosen as in Niculescu et al. ^27^: *J*_cell,background_ = 20, *J*_cell,cell_ = 100, λ_volume_ = 50, *T* = 20. The parameter λ_perimeter_ was set to 1.5, slightly lower than in Niculescu et al. ^27^ to make cells slightly more deformable. All these parameters together determine cell size, morphology, and deformability. Migratory behavior in this model is predominantly governed by two additional parameters, max_act_ and λ_act_, which were tuned as described below.

#### Phagocyte motility parameters

The phagocyte motility parameters max_act_ and λ_act_ were fitted to neutrophil motility data in a similar manner as described in “Lm motility parameters” above, now using a simulation of 300 × 400 pixels (matching the 150 × 200μm imaging window) with 40 phagocytes simulated for 1000 MCS (16min). Centroids were recorded every 5s and tracks postprocessed as described above for the Lm motility data. max_act_ and λ_act_ were tuned to match speed distributions, MSD, and autocovariance (Supp. Fig. S7), yielding max_act_ = 20 and λ_act_ = 400. In simulations with “non-motile” phagocytes, both parameters were set to 0.

#### Invasion & phagocytosis dynamics

After every step in the bacterial CPM (i.e. *v*_rel_ times per second), each bacterium was allowed to interact with the epithelium and phagocytes as follows:

1. If the bacterial pixel(s) fully overlapped a phagocyte on the phagocyte layer, it could be phagocytosed with probability *p*_ϕ_ = *k*_ϕ_/*v*_rel_; if successful, the bacterium was removed.
2. If the pixel(s) of a remaining, motile bacterium fully overlapped with a target cell on the epithelial layer, it could attach with a probability *p*_attach_ = *k*_attach_/*v*_rel_. Such bacteria can no longer leave the target cell they are attached to, but can still be phagocytosed as in (1), or can invade as in (3) below.
3. Any attached bacteria can fully invade with probability *p*_infect_ = *k*_infect_/*v*_rel”_. Once invaded, bacteria can no longer be phagocytosed. They remain in the simulation for visualization purposes but no longer affect the behavior/dynamics of any other cells.

We here assumed that attached bacteria rapidly become intracellular based on studies of endocytosis ^55,56^. We chose 5s as a starting value for initial bacteria-target cell interactions after which, bacteria become inaccessible to phagocytes during transcytosis across the epithelium. Thus, we set the average infection rate *k*_infect_ = 0.2 s^-1^. We consider the first step of attachment to be the slower, rate-limiting step with a rate *k*_attach_ = 0.051 s^-1^, consistent with kinetics from Movies S3-S6 in which Lm engaged with encountered goblets (rather than just moving past them) in about 10% of cases. The speed of phagocytosis depends on many factors but typically occurs in seconds rather than minutes ^41,57^. Because no published data exist for neutrophil phagocytosis of Lm to set the phagocytosis efficiency *k*_ϕ_ in the model, we initially set *k*_ϕ_ = *k*_attach_ = 0.051 s^-1^ for simplicity, but later investigated the sensitivity of our conclusions to both limiting rates *k*_ϕ_ and *k*_attach_.

## Supplemental Table

**Table S1:** Overview of model parameters. For details on how parameters were selected, see the Supplemental methods. Most important parameters are highlighted with •.

| Component | Parameter | Description | Value (baseline) | Further remarks |
| --- | --- | --- | --- | --- |
| GENERAL | Time resolution | # steps of the main model run for every second in real time | 1 MCS/sec | - |
| | Spatial resolution | # pixels per $\mu\text{m}$ | 2 pixels/$\mu\text{m}$ | - |
|  | Runtime | duration of the simulation | 3600 MCS (1 hour) | - |
| EPITHELIAL CPM | $V$ | # pixels per cell | 222 pixels | - |
| | $\lambda_{\text{volume}}$ | Unitless scaling of "elasticity" of the cell size | 50 | - |
| | $J_{\text{cell,background}}$ | Surface tension cell to background | 20 | - |
| | $J_{\text{cell,cell}}$ | Surface tension between epithelial cells | 30 | - |
| | $T$ | "temperature" controlling stochasticity | 20 | - |
| | $N_{\text{epi}}$ | Number of epithelial cells | $13 \times 13 = 289$ | - |
| | $N_{\text{goblet}}$ | • Number of goblet cells within $N_{\text{epi}}$ | 20 | Varied in Fig. S8 |
| BACTERIAL CPM | $\lambda_{\text{dir}}$ | • Unitless scaling of how strongly a cell "persists" in its direction of motion | 40 for Lm-RT, 0 for Lm-37 | Set to match Lm motility; see Fig. S3 |
| | $\Delta t$ | • Duration for which cells remember their previous direction, sort of "persistence time" | 60 steps in the bacterial model | $60/v_{\text{rel}} = 60/150 = 0.4\text{s}$ in real time. Set to match Lm motility; see Fig. S3 |
| | $v_{\text{rel}}$ | • # steps of the bacterial CPM run for every second in real time | 150 for Lm-RT, 0 for Lm-37 | Set to match Lm motility; see Fig. S3. Varied in Fig. S4,S10 |
| | $T$ | "temperature" controlling stochasticity | 20 | - |
| | $N_{\text{bac}}$ | • # bacteria | 100 | Varied in Fig. S8 |
| PHAGOCYTE CPM | $V$ | # pixels per cell | 314 pixels | - |
| | $P$ | Cell perimeter | 230 | - |
| | $\lambda_{\text{volume}}$ | Unitless scaling of "elasticity" of cell size | 50 | - |
| | $\lambda_{\text{perimeter}}$ | Unitless scaling of "elasticity" of perimeter | 1.5 | - |
| | $J_{\text{cell,background}}$ | Surface tension cell to background | 20 | - |
| | $J_{\text{cell,cell}}$ | Surface tension between phagocytes | 100 | - |
| | $\lambda_{\text{act}}$ | • Unitless scaling of protrusion "strength" | 400<br>(0 for non-motile) | Set to match neutrophil motility; see Fig. S7 |
| | $\text{max}_{\text{act}}$ | • Protrusion "memory" in MCS | 20<br>(0 for non-motile) | Set to match neutrophil motility; see Fig. S7 |
| | $N_{\text{ph}}$ | • # phagocytes | 14 | ~ matching the total area of goblet cells; varied in Fig. S8 |
| LAYER INTERACTIONS | $k_{\text{attach}}$ | • Rate by which motile bacteria attach to target cells | $0.051 \text{ sec}^{-1}$ | Varied in Fig. S5,S9 |
| | $k_{\text{infect}}$ | • Rate by which attached bacteria reach the "point of no return" when invading | $0.2 \text{ sec}^{-1}$ | Has a relatively small effect when $k_{\text{attach}}$ is rate-limiting |
| | $k_{\phi}$ | • Rate by which bacteria (free/attached) get phagocytosed by overlapping phagocytes | $0.051 \text{ sec}^{-1}$ | Varied in Fig. S9 |

## Supplemental Movies

Due to upload file size limitations, Movies S10-S12 are only available online. To view them, please visit: https://ingewortel.github.io/2022-listeria-goblets/.

**Movie S1: Video microscopy of Lm-37 motility in vitro**.

**Movie S2: Video microscopy of Lm-RT motility in vitro**.

**Movie S3: 2P imaging of Lm-37 interaction dynamics with mouse intestinal epithelium**.

**Movie S4: 2P imaging of Lm-RT interaction dynamics with mouse intestinal epithelium**.

**Movie S5: 2P imaging of Lm-37 interaction dynamics with human intestinal epithelium**.

**Movie S6: 2P imaging of Lm-RT interaction dynamics with human intestinal epithelium**.

**Movie S7: A CPM simulation of bacterial motility and invasion on the epithelium**. See also Fig. 4. Simulated non-motile (sLm-37) or motile (sLm-RT) bacteria, shown in blue with light-blue traces, scan the epithelium for target cells to invade (shown in a darker gray). Attached (A) and invaded (I) bacteria are shown in gray; phagocytosed (P) bacteria are invisible. Scale bar: 10 μm, timestamp in hh:mm:ss.

**Movie S8: 2D motility of neutrophils on mouse intestinal epithelium in vivo**.

**Movie S9: Motility and invasion of non-motile (sLm-37) and motile (sLm-RT) bacteria on the epithelium in the presence of phagocytes**. The epithelium is shown in gray, with darker cells representing target cells. Scanning bacteria are shown in blue with their traces in lighter blue; attached (A) and invaded (I) bacteria are shown in gray, phagocytosed (P) bacteria are no longer shown. Phagocytes are shown as pink cells with dark protruding fronts. Scale bar: 10 μm, timestamp in hh:mm:ss. See also Movie S7.

**Movie S10: Motility and invasion of non-motile (sLm-37) and motile (sLm-RT) bacteria for varying challenge doses**. As Movie S9, but now for varying numbers of bacteria. Scale bar: 10 μm, timestamp in hh:mm:ss. A = attached bacteria, I = invaded bacteria, P = phagocytosed bacteria.

**Movie S11: Motility and invasion of non-motile (sLm-37) and motile (sLm-RT) bacteria for varying numbers of target cells**. As Movie S9, but now for varying numbers of target cells. Scale bar: 10 μm, timestamp in hh:mm:ss. A = attached bacteria, I = invaded bacteria, P = phagocytosed bacteria.

**Movie S12: Motility and invasion of non-motile (sLm-37) and motile (sLm-RT) bacteria for varying numbers of phagocytes**. As Movie S9, but now for varying numbers of phagocytes. Scale bar: 10 μm, timestamp in hh:mm:ss. A = attached bacteria, I = invaded bacteria, P = phagocytosed bacteria.

## Supplemental Figures

**Figure S1:**
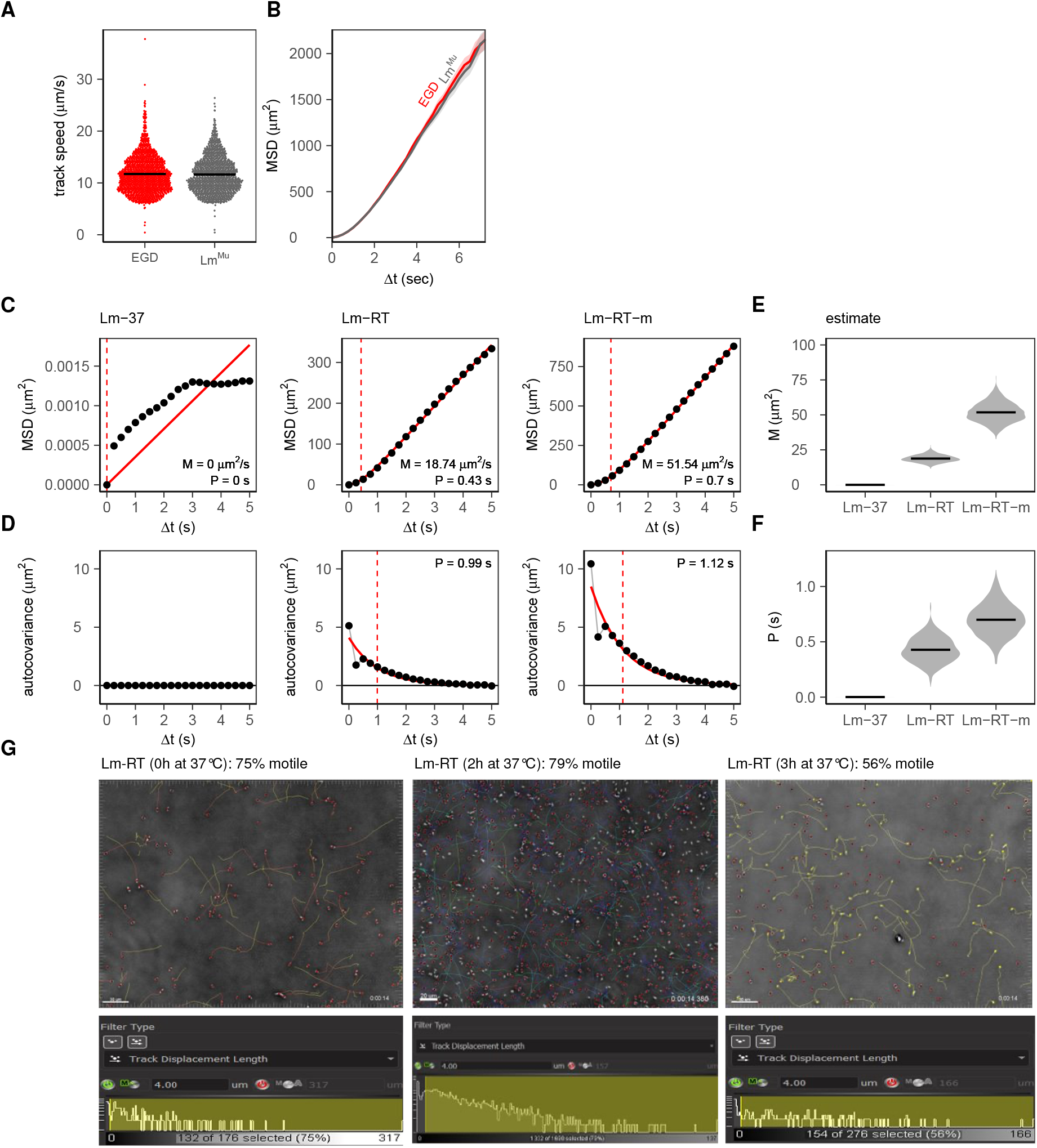
Comparing Lm motility across populations. **(A**,**B)** Comparison of speed (A) and mean squared displacement (MSD, B) between human EGD Lm and murinized Lm^Mu^. **(C)** MSD curve of the three Listeria populations (EGD Lm-37, Lm-RT, Lm-RT-m), fitted by Fü rth’s equation (red line). The red dashed line indicates the persistence time as determined from the fit. Curves were fit on Δt up to 5s; for longer Δt, fast cells tend to leave the imaging window and the MSD becomes biased (see methods for details). Lm-37 did not move and could not be fitted by Fü rth’s equation. **(D)** Autocovariance curve of the populations as fitted by an exponential decay: *f* (*x*) = *f*_0_ ∗ exp(−*x*/*P*), red line. This yields a slightly higher estimate of the persistence time P (vertical dashed red lines), but still in the same order of magnitude as those in panel C. **(E**,**F)**, To estimate uncertainty in motility parameters estimated from the MSD (motility coefficient M and persistence time P), tracks were resampled from the original populations with replacement N = 1000 times, to obtain N “bootstrapped” datasets of equal size as the original. Resampled datasets were then fitted with Fü rth’s equation as shown in panel A to obtain N estimates of M and P. **(G)** To assess how long Lm-RT stay motile after being placed at 37°C, Lm-InlA was first grown at RT with shaking (200rpm) to OD600 around 1.0, and then switched to a 37°C incubator (shaking at 200rpm) to assess motility after 2-3 hours. Samples were diluted 1:10 with BHI and plated on a non-charged slide (Globe Scientific Cat No 1324W); a 2D time lapse video was recorded for 15 seconds with 250ms time and 100ms exposure using Olympus IX51 inverted microscope with 20X objective and phase dichroic filter. Cells were were tracked in Imaris 9.3 and % motile cells were calculated using 4 μm track displacement length filter.

**Figure S2:**
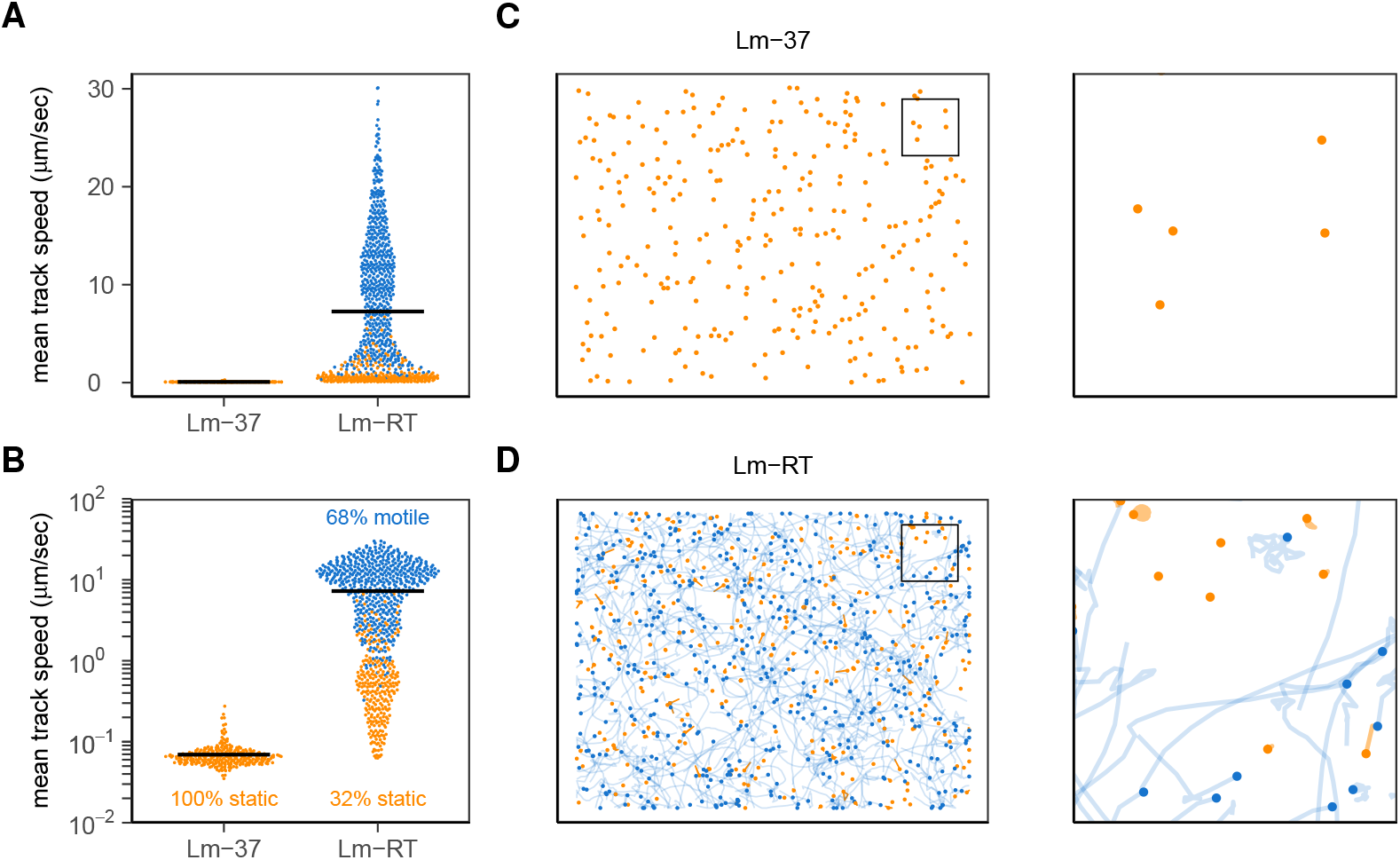
Removing non-motile cells from the Lm-RT population to create a more representative motile data set. To remove artefacts of non-motile cells sticking to the glass slide, bacteria separate tracks were separated into “motile” vs “static” tracks. Briefly, tracks were considered motile whenever the track coordinates were described better by two Gaussian distributions (splitting the track in two parts) than by a single Gaussian. If a single Gaussian distribution was a reasonably good fit for the observed coordinate, the tracks were classified as static. See Supplemental methods for details. **A**,**B**, static (orange) and motile (blue) cells of Lm-37 and Lm-RT, shown in the speed distribution. Whereas static tracks tend to have low speeds, there are also some static tracks with relatively high speeds (mostly when the track contains a single motile step while the cell otherwise does not move). Blue tracks represent the “Lm-RT-m” population. **C**,**D** tracks of Lm-37 and Lm-RT, showing that the filter indeed reasonably removes non-motile cells. Zoomed inset: 50 × 50 μm.

**Figure S3:**
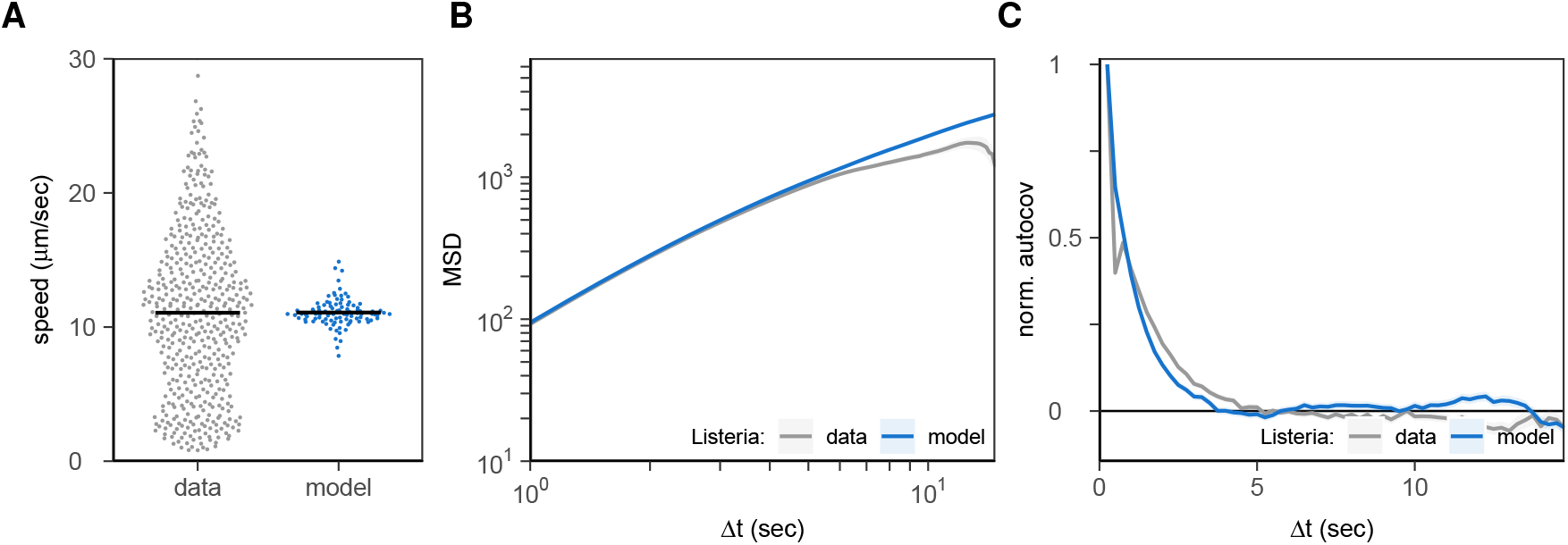
Matching motility of simulated Lm to in vitro data. Average bacterial motility (of 50 simulated bacteria) in the model (sLm-RT) closely matches in vitro motility (Lm-RT) in both speed and directionality, as shown by: **A**, the distribution of cell speeds, **B**, the mean squared displacement (MSD) over different time intervals Δt and **C**, the (normalized) autocovariance of movement “step” vectors with time Δt between them (the longer it takes for this curve to drop to zero, the larger the persistence time of the cells). See Methods for details.

**Figure S4:**
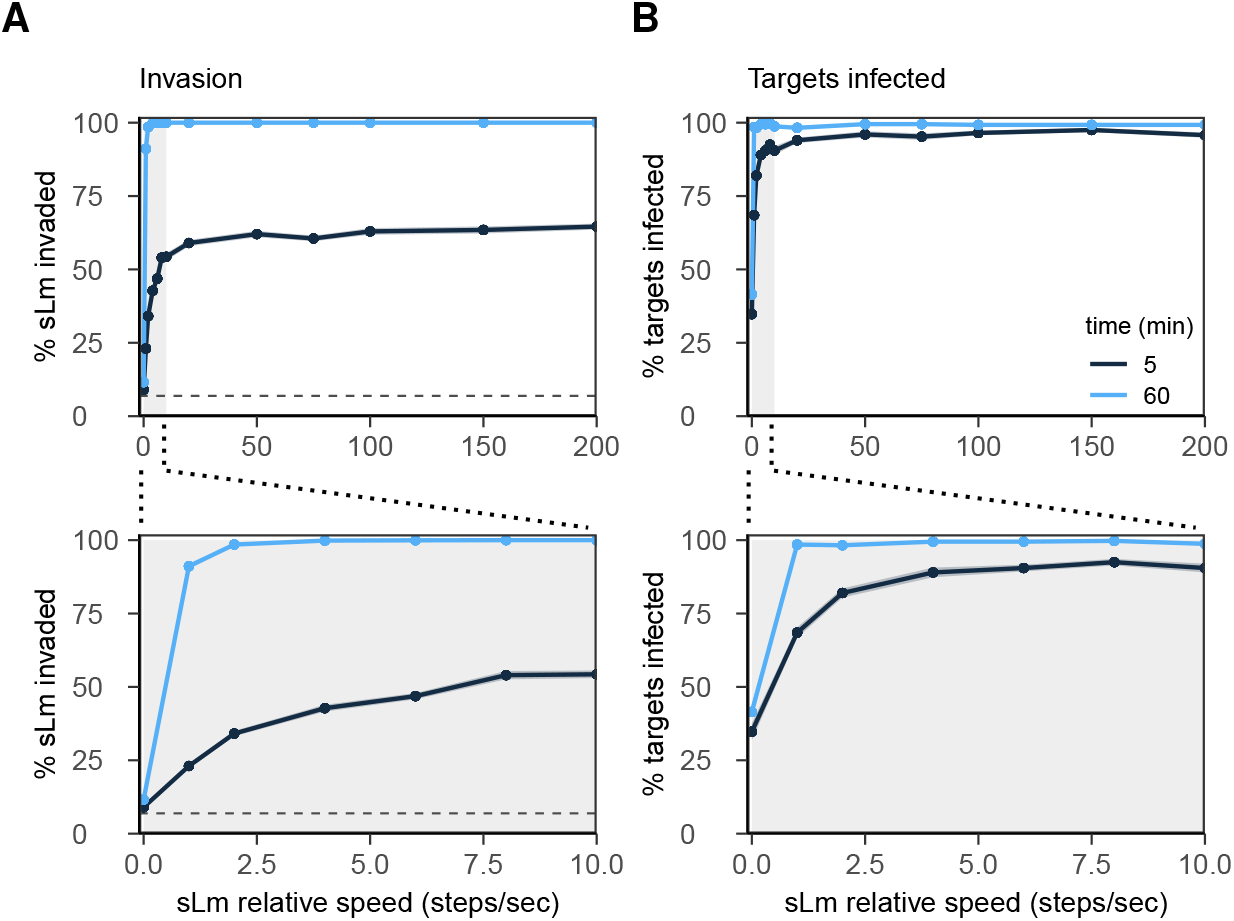
Target cell infection efficiency as a function of sLm speed. sLm speed was changed by varying the v_rel_ parameter in the model. This parameter controls how many “steps” of the bacterial model occur each second; high values increase bacterial speed whereas a value of zero means that bacteria are completely static (the default value used throughout the paper is 150 to simulate sLm-RT and 1 to simulate sLm-37). Target cell infection efficiency is measured as **A**, the % of sLm that have invaded after 60 min for all tested speeds (top) or for sLm speeds up to 10 steps/s (bottom; corresponding to the gray region in the upper plot); and **B**, likewise, but now for the % of target cells that has been invaded by sLm. Results are shown as mean±SE for 20 independent simulations. Horizontal dashed lines in **A** represent the % of the surface area covered with target cells.

**Figure S5:**
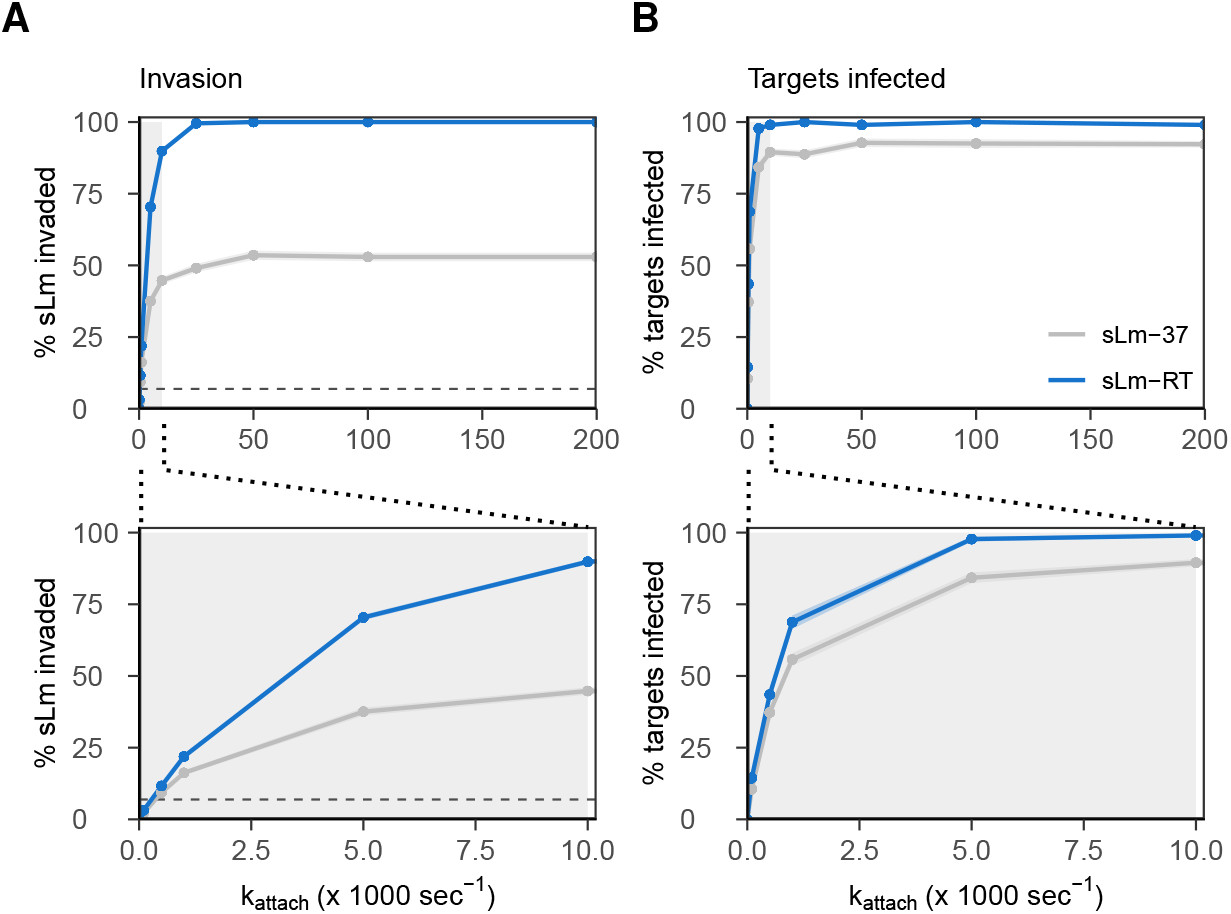
Target infection efficiency as a function of the attachment rate k_attach_ of sLm to target cells. When k_attach_ is high, scanning the epithelium will immediately attach to any target cell they encounter; when it is low, sLm are more likely to move past target cells instead of attaching to them (default value used in the paper: 0.051 s^-1^). Target infection efficiency is measured as **A**, the % of sLm that has invaded after 60 min for all tested speeds (top) or for sLm speeds up to 10 steps/s (bottom; corresponding to the gray region in the upper plot); and **B**, likewise, but now for the % of target cells that has been invaded by sLm. Results are shown as mean±SE for 20 independent simulations.

**Figure S6:**
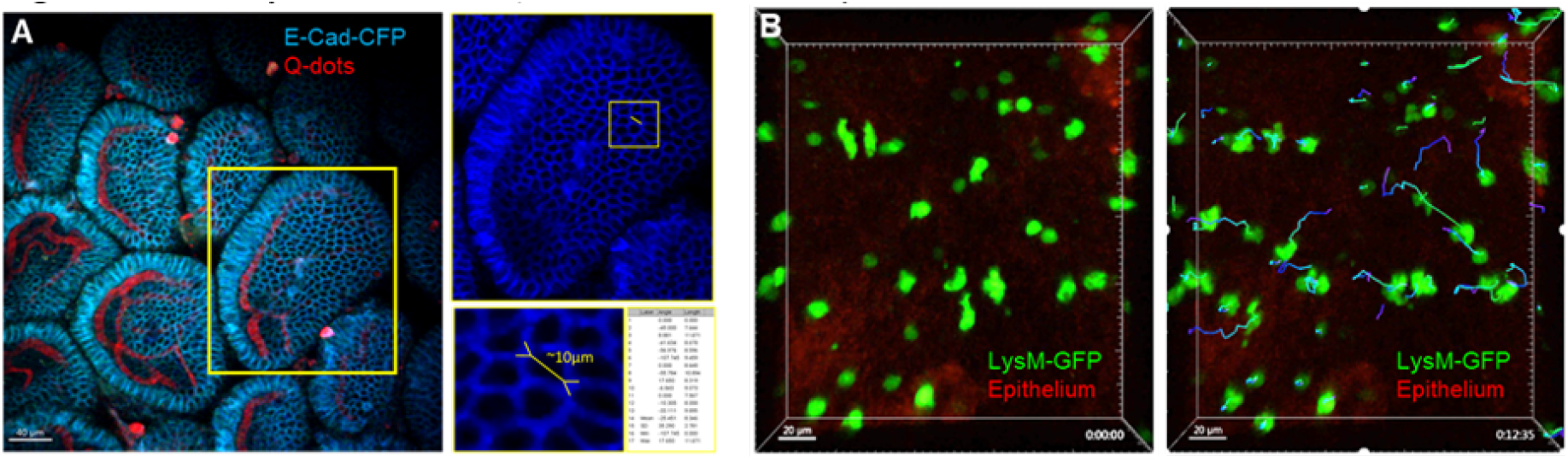
2D motility of neutrophils on mouse intestinal epithelium in vivo.

**Figure S7:**
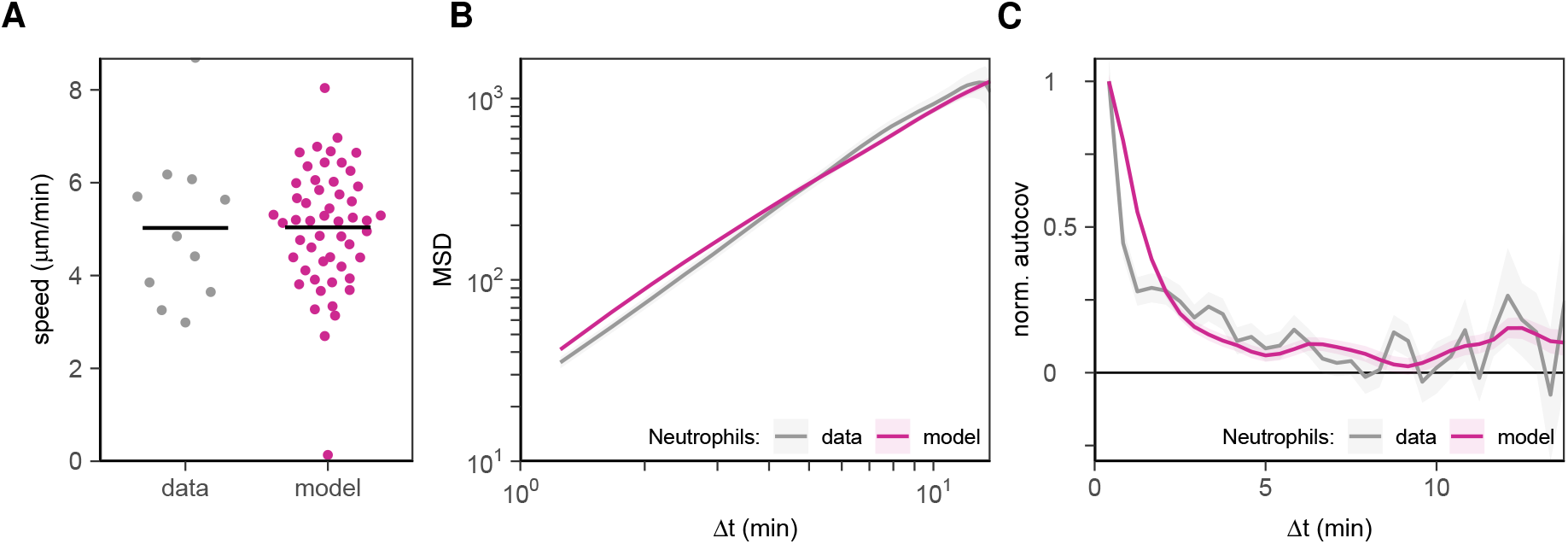
Matching simulated phagocyte motility to in vivo neutrophil motility. Phagocyte motility in the model (average of 40 simulated cells) closely matches in vivo motility of neutrophils crawling between epithelium and coverglass. Motility matches in both speed and directionality, as shown by: **A**, the distribution of cell speeds, **B**, the mean squared displacement (MSD) over different time intervals Δt and **C**, the (normalized) autocovariance of movement “step” vectors with time Δt between them (the longer it takes for this curve to drop to zero, the larger the persistence time of the cells). See Methods for details.

**Figure S8:**
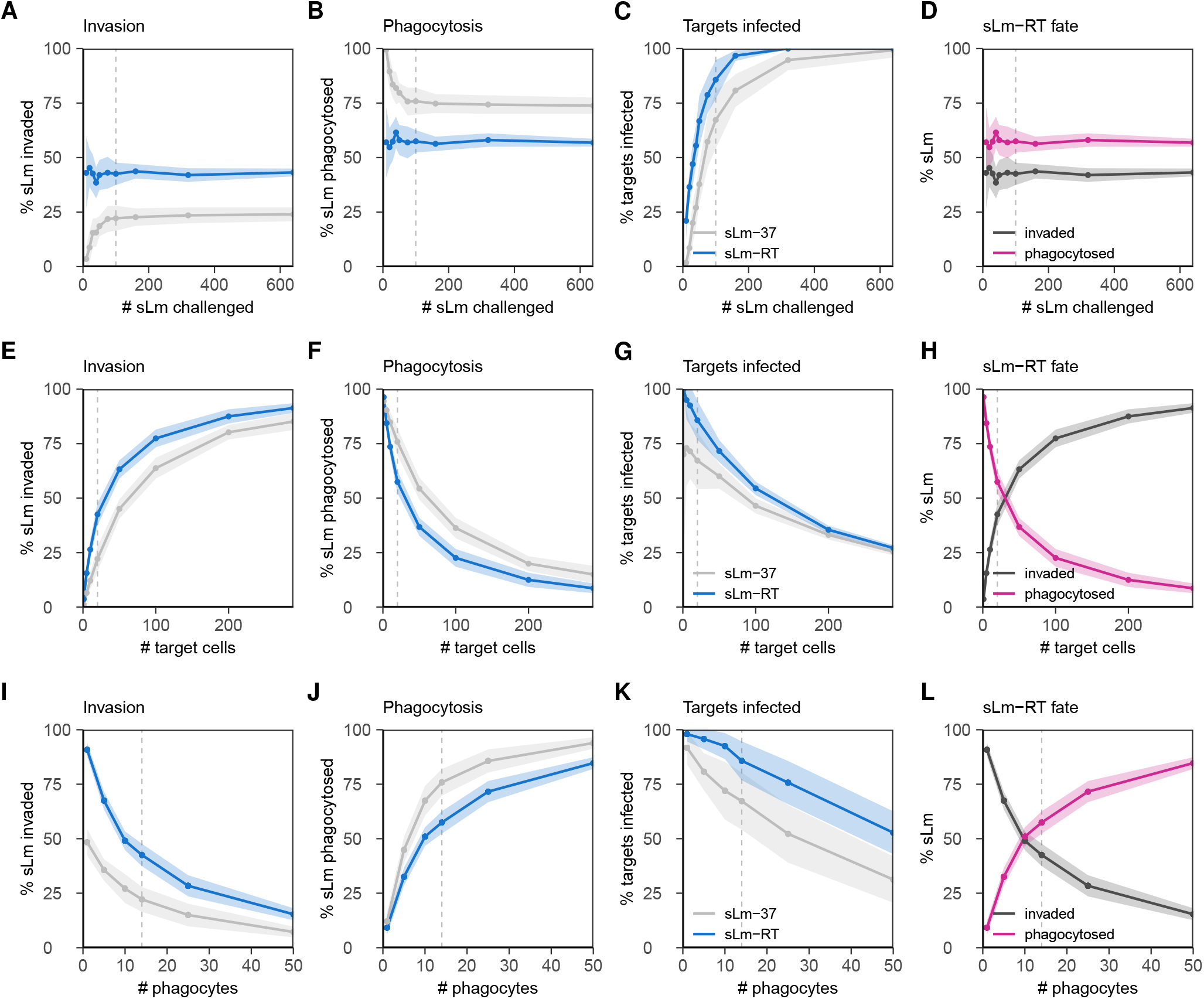
Dependency of immunological outcomes on numbers of bacteria, target cells, and phagocytes. Immunological outcomes were assessed after varying **A-D**, the number of sLm challenged (with **A**, % sLm invaded, **B**, % sLm phagocytosed, **C**, % of target cells invaded by sLm after 60 min, and **D**, the data from panels **A-B** compared in one plot for motile sLm-RT). While challenge dose does not affect the % of sLm invading or phagocytosed for motile sLm-RT (blue), it does for low doses of the non-motile sLm-37 (gray), which are more likely to diffuse on or near target cells before being phagocytosed. Dependency of immunological outcomes was similarly tested for **E-H**, the number of “target” cells (i.e. goblets that can be invaded; the total number of epithelial cells is 289), and **I-L**, the number of phagocytes. Results are shown as mean ±standard deviation (SD) for 20 independent simulations for motile (sLm-RT, blue) and non-motile bacteria (sLm-37, gray). Vertical dotted lines indicate the baseline of 100 sLm, 20 target cells, and 14 phagocytes used in the rest of the paper.

**Figure S9:**
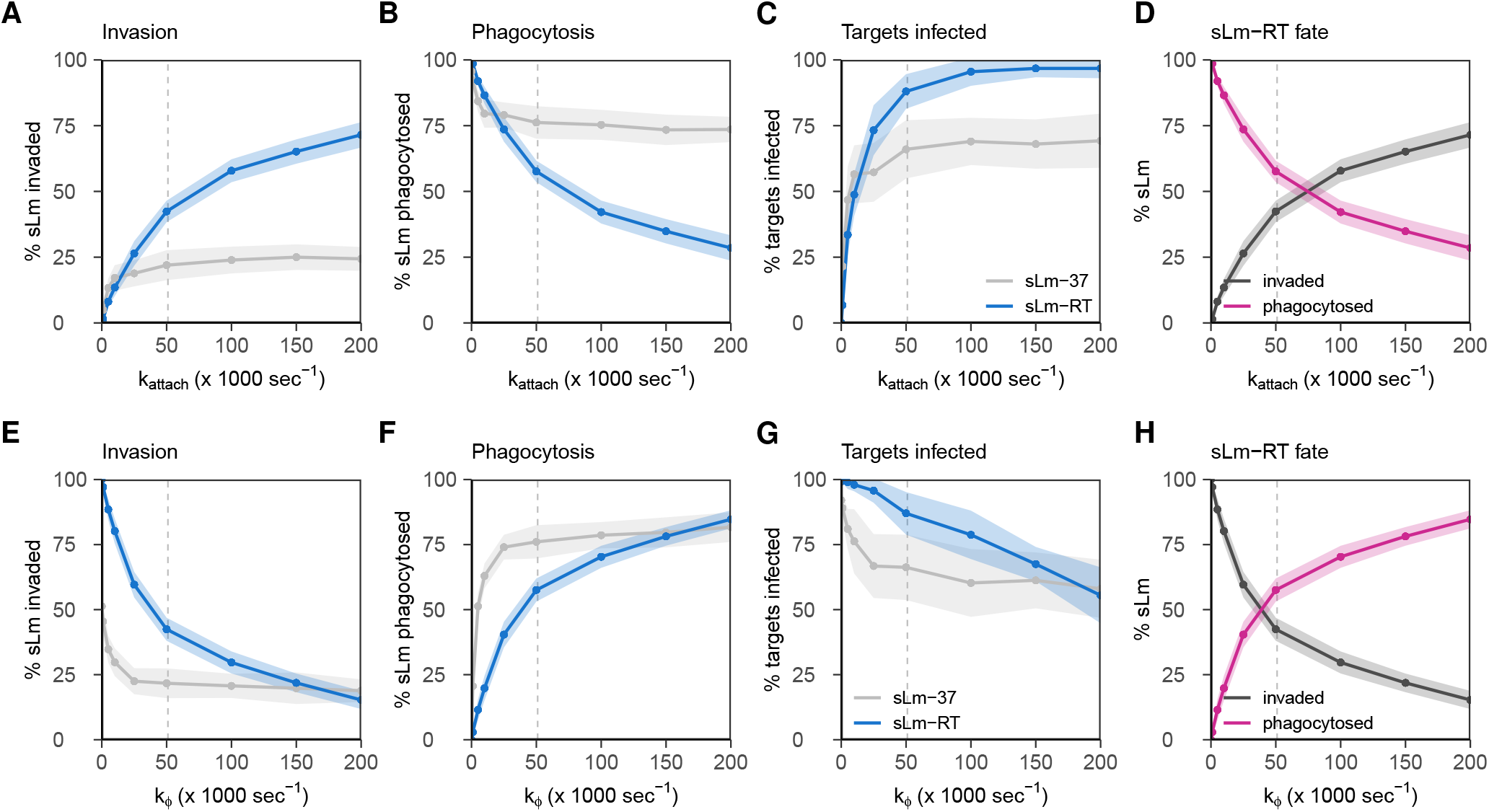
Dependency of immunological outcomes on invasion and phagocytosis kinetics. Immunological outcomes were assessed after varying **A-D**, the rate by which sLm attach to target cells they encounter (k_attach_) or **E-G**, the rate by which sLm are phagocytosed by encountered phagocytes (k_φ_). Plots represent the following: **A**,**E**, % sLm invaded, **B**,**F**, % sLm phagocytosed, **C**,**G**, % of target cells invaded after 60 min, and **D**,**H**, the data from panels A-B compared in one plot for motile sLm-RT. Results are shown as mean±SD for 20 independent simulations for motile sLm-RT (blue) and non-motile sLm-37 (gray). Vertical dotted lines indicate the default values of k_attach_ = k_φ_ = 0.051 s^-1^ used in the rest of the paper.

**Figure S10:**
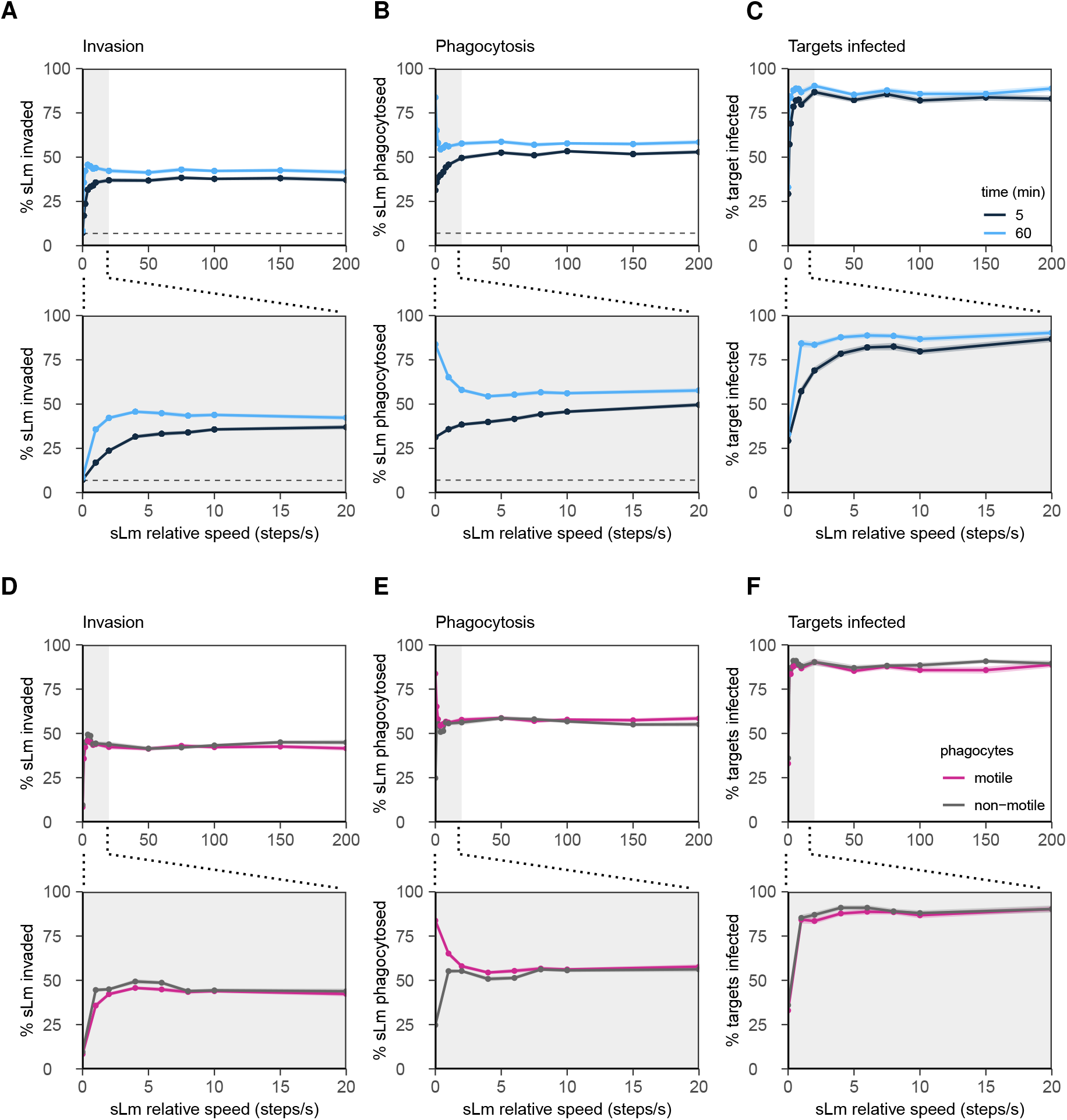
Dependency of immunological outcomes on sLm and phagocyte speed. Immunological outcomes were assessed after varying **A-C**, the relative sLm speed in steps/s (see also Supp. Fig. S4), and **D-F**, phagocyte speed. Plots show A, % invaded sLm, B, % phagocytosed sLm and C, % target cells invaded after 5 or 60 min, with horizontal lines in **A-B** indicating the percentage of the surface area covered by target cells or phagocytes, respectively. The bottom plots in **A-C** are zoomed in on lower relative speeds, indicated by the gray shaded region in the top-row panels. **D-F** show only the 60-minute curve, but now for motile phagocytes (modelled with default parameters, pink) compared to non-motile phagocytes (modelled with λ_act_ = max_act_ = 0, gray). Phagocyte motility affects outcomes only in the (very) low range of sLm motility, where bacteria are static enough that phagocytes can actually “hunt” them. All lines represent mean ±SE of 20 independent simulations.

